# A population of stress-like cancer cells in melanoma promotes tumorigenesis and confers drug resistance

**DOI:** 10.1101/396622

**Authors:** Maayan Baron, Mohita Tagore, Miranda V. Hunter, Isabella S. Kim, Reuben Moncada, Yun Yan, Nathaniel R. Campbell, Richard M. White, Itai Yanai

## Abstract

Transcriptional profiling has revealed a diverse range of cancer cell states, however an understanding of their function has remained elusive. Using a combination of zebrafish melanoma modeling and human validation, we have identified a conserved stress-like state that confers intrinsic drug resistance. The stress-like state expresses genes such as *fos*, *hsp70* and *ubb*, all required for adaptation to diverse cellular stresses, and we confirmed its existence using immunofluorescence and spatial transcriptomics. We provide evidence that this state has a higher tumor seeding capabilities compared to non-stressed cells, and confers intrinsic resistance to MEK inhibitors, a commonly used melanoma therapeutic. Furthermore, the stress-like program can be induced by extrinsic processes such as heat shock, and confers resistance to both MEK and BRAF inhibitors in both zebrafish and human melanomas. Collectively, our study suggests that the transcriptional states associated with therapeutic failure are established during the earliest steps of tumorigenesis.

## INTRODUCTION

A universal feature of cancer is its genetic and phenotypic heterogeneity (Fisher et al., 2013; Lawrence et al., 2013; Meacham and Morrison, 2013; Sharma et al., 2010). Genetically, tumor evolution leads to a recurring set of DNA alterations in genes such as KRAS, BRAF or p53. (Hollstein et al., 1991; Lièvre et al., 2006; Riely et al., 2009; Vogelstein and Kinzler, 2004). In addition to alterations of DNA, transcriptional heterogeneity of cancer cells is increasingly recognized in a diverse array of tumors including melanoma (Jerby-Arnon et al., 2018; Rambow et al., 2018; Tirosh et al., 2016a), glioblastoma (Patel et al., 2014), oligodendroglioma (Tirosh et al., 2016b), breast (Kim et al., 2018) and head and neck cancer (Puram et al., 2017, 2018). In glioblastoma, multiple transcriptional programs of cancer cells co-exist according to classical, proneural, neural, and mesenchymal cell states (Patel et al., 2014). For melanoma, the genetics have been well characterized in terms of recurring genes and pathways across nevus and invasive bulk tumors (Cancer Genome Atlas Network, 2015; Tsao et al., 2012), and recent work using single-cell RNA sequencing (scRNA-Seq) has mapped out multiple transcriptional programs (Jerby-Arnon et al., 2018; Rambow et al., 2018; Tirosh et al., 2016a). However, despite the characterization of this diversity, the functional consequences of cancer cell states are not well understood.

Much like genetic evolution during tumorigenesis, transcriptional evolution can give rise to different cancer cell states over time. Many open questions remain about the nature of these cell states, including how they arise, their physical organization, and their functional consequences (Barkley and Yanai, 2019). In addition, evidence has been presented indicating that cell states are not fixed, which may be an important aspect of tumor cell plasticity (Hoek et al., 2008; Verfaillie et al., 2015; Widmer et al., 2012). Switching between states can be driven by factors from the microenvironment such as WNT5A (O’Connell et al., 2013) and EDN3 (Kim et al., 2017), or from varying levels of MITF (Carreira et al., 2006; Vivas-García et al., 2019), the master transcription factor for melanocyte development. The ability of individual cells to take on these varying transcriptional states also has important consequences for patient prognosis (Sarrió et al., 2008). For example, the acquisition of an AXL^high^/MITF^low^ state is associated with an invasive, metastatic phenotype (Müller et al., 2014; Tirosh et al., 2016a), and phenotype switching has been associated with drug tolerance (Ahmed and Haass, 2018). More recently, a transcriptional state associated with RXR signaling has been found to be a key factor in resistance to BRAF/MEK inhibitors (Rambow et al., 2018).

To study the origins and function of these various cell states a system is required for the prospective identification and manipulation of these cell states within an intact microenvironment. To address this, we have developed a transgenic zebrafish model of melanoma, in which the *BRAF^V600E^* oncogene is expressed in melanocytes (Ceol et al., 2011; Patton et al., 2005; White et al., 2011). These fish develop melanomas that resemble the human disease at histological, transcriptomic and genomic level, and have previously been shown to be a powerful model for the study of both metastasis and drug resistance (Heilmann et al., 2015; Patton et al., 2005; Yen et al., 2013). Importantly, the tumors can be repeatedly sampled over time, something that is difficult to do in humans. This allows us to understand when these cell states arise over the course of tumor evolution.

Previous scRNA-Seq melanoma studies have revealed multiple cell states in melanoma, including neural crest, pigmented, invasive and starved states (Rambow et al., 2018). We and others have previously shown that the neural crest transcriptional program, typified by genes such as *SOX10*, is essential to melanoma initiation because it provides the proper mileu on which DNA mutations can act (Kaufman et al., 2016; Shakhova et al., 2012; Travnickova et al., 2019; White et al., 2011). In contrast, the existence of other cell states such as a “stress-like” state, expressing genes such as *fos* and *jun*, has been suggested by prior work but its functional role remains unclear (Tirosh et al., 2016a). Further complicating the matter is the observation that a stress transcriptional program has been shown to arise as an artifact of cell dissociation protocols and flow cytometry sorting (van den Brink et al., 2017). This study highlights the challenge of identifying a role for stress signaling in cancer, which is likely biologically meaningful but can be strongly affected by artifacts of single-cell methods. Collectively, the existence of the stress-like state as a bona fide biological property of tumorigenesis remains unclear.

Here we use a zebrafish model of melanoma to study stress-like cancer subpopulations. Using spatial transcriptomics on intact tumors, which does not rely on cell sorting or dissociation, we show that the stress-like program is a bona fide cell population. Importantly, this stress-like program is found across a wide variety of tumor types, including triple-negative breast cancer, oligodendroglioma and pancreatic adenocarcinoma, suggesting a conserved feature of cancer. We developed a transgenic reporter of this stress-like state, and used it to demonstrate that this population is intrinsically drug resistant, and arises early in tumor evolution. Our work has implications for the treatment of human cancer, since it implies that the transcriptional states associated with therapeutic failure - much like genetic forms of resistance - are likely built into the origins of the tumor.

## RESULTS

### Cell state mapping reveals conserved cell types between zebrafish and human melanomas

Animal models of cancer allow for both characterization and functional analysis of cell states in cancer. To study this in melanoma, we utilized a transgenic zebrafish model in which human *BRAF^V600E^* is expressed in melanocytes through the *mitfa* promoter. When expressed in a *p53*-/- background, these fish develop melanomas that resemble the human tumors (Patton et al., 2005). To determine if these tumors recapitulate the cellular diversity of human melanoma, we performed scRNA-seq on n=8 tumor biopsies from these transgenic tumors. We processed ∼15k cells from the eight samples (Figure 1a), using the inDrop system (Klein et al., 2015) to comprehensively capture the cell types present in the tumor. After quality control and filtering (see Methods), we were left with a total 10,012 cells, each with an average of approximately 2,500 transcripts and 1,000 genes detected (Supplementary Figure 1). Studying the transcriptomes, we detected eight cell types, each forming a distinct cluster when visualized using tSNE (t-distributed stochastic neighbor embedding) (Maaten and Hinton, 2008) (Figure 1b). To annotate these clusters, we first identified the cancer cell population by the detection of *BRAF^V600E^* transcripts. Known markers were used to annotate the rest of the cell types: keratinocytes, fibroblasts, erythrocytes, natural killer cells, neutrophils, macrophages, other lymphocytes (Supplementary Table 1). For example, a cluster was annotated as keratinocytes since it was enriched in the expression of keratin 4 (krt4) and other genes. In humans, keratinocytes are the major cell type that surrounds the melanocytes from which these tumors arise, and contribute important growth factors such as EDN3 that promote tumor growth (Saldana-Caboverde and Kos, 2010). Other identified cell types include immune cells such as T-cells and macrophages, also commonly observed cell types in the human disease. These data confirm that the zebrafish melanomas are composed of multiple cell types that closely resemble the cell types seen in humans.

**Figure 1.**
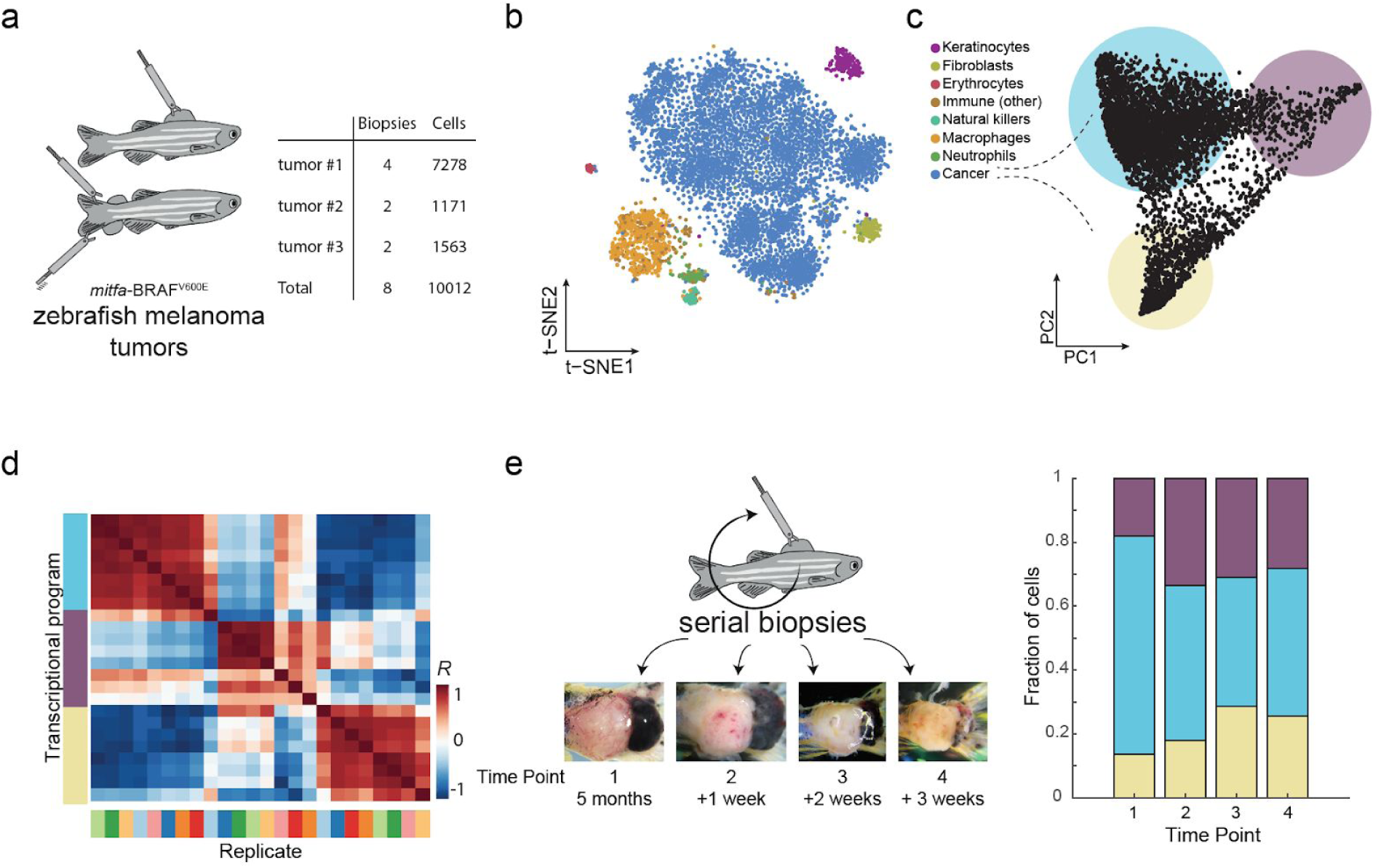
Single-cell RNA-Seq on zebrafish melanoma. (a) Eight tumor biopsies were processed from three distinct tumors using scRNA-Seq. (b) tSNE analysis of 7,278 single cells from tumor #1. Color indicates the inferred cell type. (c) PCA on the tumor cancer cells revealed three transcriptional cell states, indicated by the colored circles. (d) Heatmap showing the Pearson’s correlation coefficients between the three cell states across all eight biopsies. Biopsy samples cluster according to states and not tumor or animal of origin. (e) Serial biopsies were taken from the same tumor (tumor #1) in one week intervals. Micrographs show the tumor at each time point. The bar plot indicates the proportions of the cell states detected in panel (c) for each biopsy.

### Melanomas have a cell stress-like transcriptional program as seen by scRNA-seq

To study transcriptional heterogeneity within the melanoma cell populations, we applied principal component analysis (PCA) on only the cells expressing the human *BRAF^V600E^* gene. This analysis revealed a triangle-shaped arrangement of transcriptomes with a concentration of cells near the vertices (Figure 1c). We verified that this shape is not driven by pre-processing methods or any technical aspects such as number of UMI detected (Supplementary Figure 2). Furthermore, analyzing the cells across our eight biopsies, we found that all three transcriptional states are present in each sample (Figure 1d). In order to test whether these 3 cancer cell states are present in early stages of tumorigenesis, we performed microscopic biopsies on one of the transgenic zebrafish tumors (tumor #1) as soon as they were visible on the skin of the animals followed by scRNA-Seq. We found that all three states were readily identifiable as early as 5 months after tumor initiation and that their proportion within each biopsy does not change significantly over time (Figure 1e). These results support the notion that melanoma cancer cells express at least three distinct transcriptional states from early stages of tumorigenesis.

To characterize the functional attributes of the three cancer cell states, we studied their underlying transcriptional programs in terms of uniquely expressed genes (Figure 2a). For this we performed differential gene expression analysis among the groups of 500 cells closest to each PCA vertex in Figure 1c. We found that transcriptional program 1 (low PC1, low PC2) is enriched for the expression of neural crest genes, such as *sox2* and *sox10*, suggesting co-option of this progenitor program by the cells (Figure 2b). In contrast, transcriptional program 2 (low PC1, high PC2) is enriched with the expression of genes associated with mature melanocytes, such as *dct*, *tyrp1b*, and *pmela*, indicating that these cells co-opt the differentiated melanocyte transcriptional program (Figure 2b). To test for the distinction between program 1 and 2, we compared their expression profiles to a published dataset that measured gene expression changes that accompany human melanocyte differentiation from pluripotent stem cells (Mica et al., 2013). As expected, we found that programs 1 and 2 best correlate with the expression to neural crest cells and mature melanocytes, respectively (Figure 2c), further supporting the notion that these states capture distinct stages of melanocyte differentiation process.

**Figure 2.**
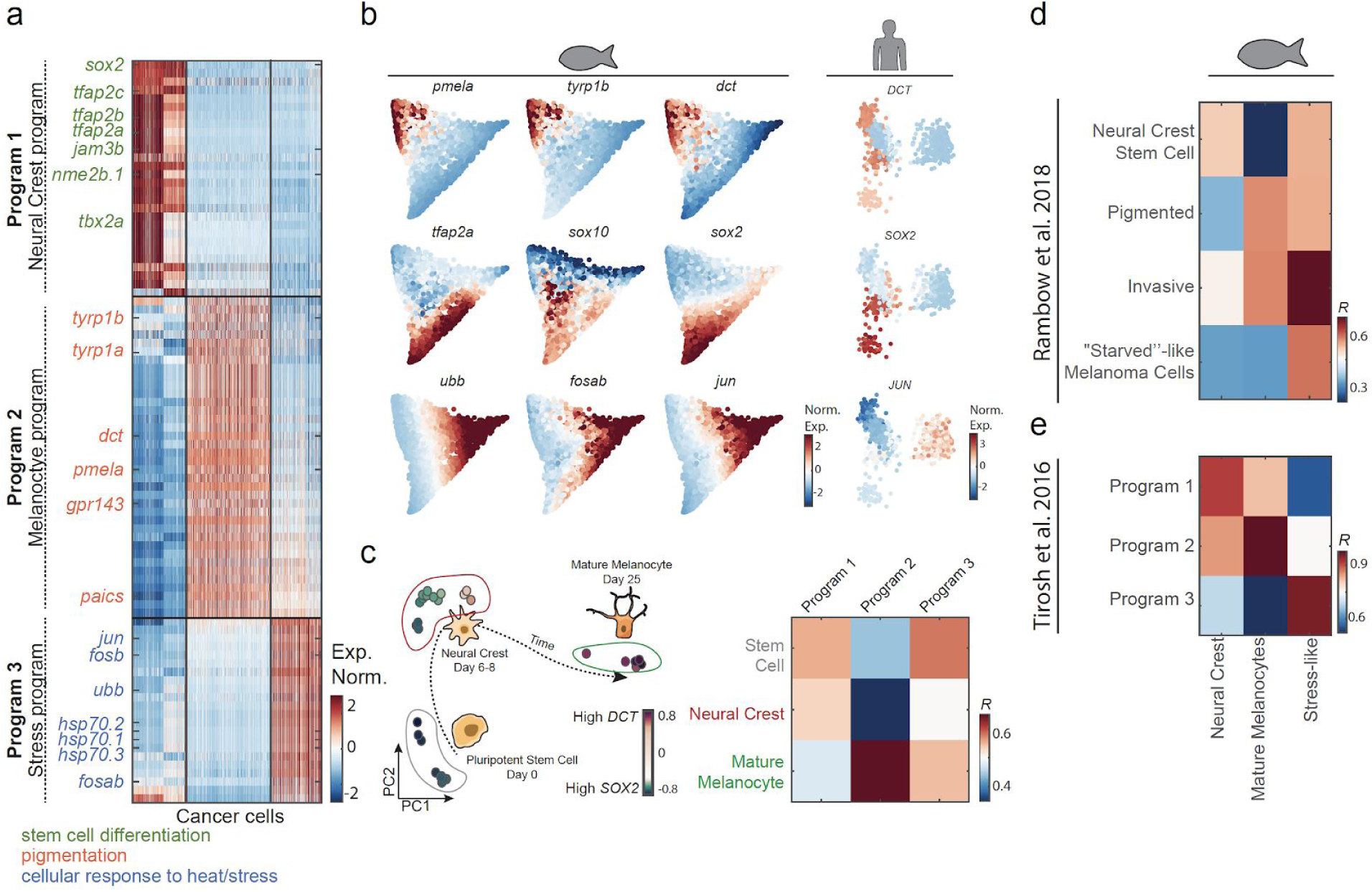
Transcriptional program underlying cancer cell states. (a) Normalized expression levels of the differentially expressed genes across the cancer cells of tumor #1. Genes are colored by function based on GO annotations indicated at the bottom. (b) Gene expression levels for the indicated genes, mapped onto the PCA shown in figure 1c (left three columns) and PCA from previously reported human scRNA-seq data-set (Tirosh et al., 2016a). (c) PCA of bulk melanocyte differentiation from previously reported data (Mica et al., 2013), colors indicate expression levels of *SOX2* (purple) and *DCT* (green). In the heatmap, the Pearson’s correlation levels are shown between the three human melanoma cell type programs and the developmental transcriptomes of stem cells, neural crest, and mature melanocytes. (d,e) Comparison between zebrafish melanoma transcriptional program and (d) human melanoma transcriptional program (Tirosh et al., 2016a) and (e) patient-derived xenografts (Rambow et al., 2018).

We found that transcriptional program 3 is enriched in the expression of genes such as *jun*, *fosb*, *fosab*, *ubb*, and heat-shock response genes, all associated with a stress-like transcriptional program. This was of interest to us since these classes of genes were previously found to be a potential artifact of scRNA-Seq methods (van den Brink et al., 2017), but yet were confined to only the cancer cells in our study despite the fact that all cells were treated the same way. Together, this analysis led us to conclude that the zebrafish melanomas contain at least three different cell states that co-opt existing transcriptional programs corresponding to neural-crest cells, mature melanocytes, and a stress-like response, which are consistent across our eight biopsies (Supplementary Figure 3).

To address whether our identified cancer cell states are unique to the zebrafish system or conserved also in human melanoma, we compared them to the cell states detected in two previously reported human melanoma scRNA-Seq datasets. We first compared our data to that of Rambow et al. (Rambow et al., 2018), who described four transcriptional states of melanoma cells isolated from patient-derived xenografts that were exposed to RAF/MEK inhibition. Comparing our three cell states to the four identified by this study, we found that the neural crest state, the mature melanocyte, and the stress-like state showed the highest respective gene expression correlation with Rambow et al.’s annotated neural crest stem cells, pigmented cells, and ‘starved’-like melanoma cells respectively (Figure 2d). We next compared our data to the Tirosh et al. scRNA-seq data on a panel of human melanoma samples. Focusing on the MITF population since the zebrafish *BRAF^V600E^* transgene is driven by the *mitfa* promoter, we found that these have a similar pattern of expression to the zebrafish tumor expression (Figure 2b,d). Overall, these analyses suggest that our observations of zebrafish cancer cell states constitute a conserved aspect of human melanoma.

### Spatial transcriptomics supports presence of stress-like cancer cells

Since the neural crest and melanocyte programs are well described in melanoma, we focused on the poorly characterized stress-like state. Since it has been observed that a stress transcriptional program may arise as an artifact of cell dissociation protocols and flow cytometry sorting (van den Brink et al., 2017), the existence of the stress-like state as a bona fide biological property has been called into question. Supporting the existence of a stress-like cell state, we found that the associated transcriptional program was not detected in two other non-cancer datasets generated using the same dissociation and scRNA-Seq methods (Supplementary Figure 4). In addition, the expression levels of the stress-like program are significantly lower in the non-cancer datasets as compared to the melanoma dataset (Supplementary Figure 4). We also implemented the previously described *in silico* purification method (van den Brink et al., 2017) to our dataset, and found that the stress-like state remains across a range of thresholds (Supplementary Figure 4).

To more directly validate the existence of this stress-like cell state in melanoma using an orthogonal technology, we turned to spatial transcriptomics (Ståhl et al., 2016). This is an *in situ* RNA-sequencing approach that does not depend upon dissociation or flow sorting of cells. From a transgenic zebrafish melanoma, we derived a stable cell line called ZMEL1-GFP (Heilmann et al., 2015), and then transplanted these into recipient transparent *casper* zebrafish, to aid in visualization of the tumor. After 2 weeks, the recipient animals formed large, GFP-positive melanomas. We prepared frozen sections of the tumor and surrounding normal tissue, along with several sections from non-tumor bearing regions of the fish. Each section was placed on a spatial transcriptomics slide containing spatially barcoded mRNA probes, allowing us to capture the mRNA from the section and prepare cDNA libraries with added spatial information. We then analyzed this data for all 3 of the melanoma cell states we had observed in our initial scRNA-Seq experiment. Comparing this data to a hematoxylin/eosin stain of the same section enables the annotation of tumor and normal areas. This confirmed that the expression of genes such as *sox2* (neural crest), *pmela* (melanocyte) and *fosab* (stress) are highly enriched in the tumor area when compared to the normal surrounding tissue (Figure 3a-b, *P*<10^−7^, Mann-Whitney test, Supplementary Figure 5). Extending this analysis to all genes for each of the 3 cancer-cell states, we found that overall all three states are highly enriched in the tumor area (Figure 3c and Supplementary Figure 5) when compared to surrounding normal tissue. As a negative control, we examined randomly selected genes (n=200), and found that these do not show an enrichment in the tumor. In addition, we also found that the expression of the genes of each of the cancer cell states were not found in non-tumor bearing sections of animals (Supplementary Figure 5). Collectively, these data support our findings that the 3 observed cancer cell states in our tumors - neural crest, melanocyte and stress-like - are not an artifact of cell dissociation and sorting and represent actual states in intact tissues.

**Figure 3.**
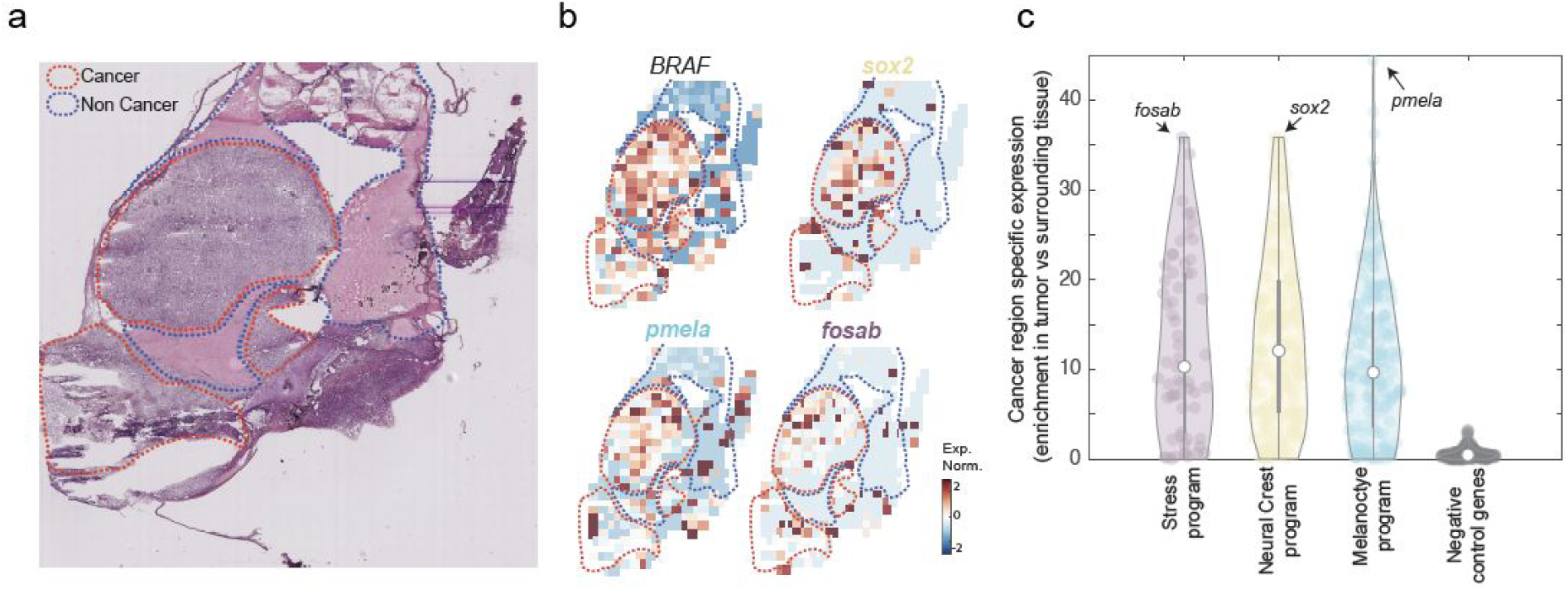
The transcriptional programs of the cancer cell states are enriched in cancer areas. (a) Hematoxylin and eosin stain of a zebrafish transplanted tumor section. Red and blue dotted lines mark cancer and non-cancer areas, respectively. (b) Gene expression profiles of the indicated genes obtained by spatial transcriptomics performed on a section adjacent to the one shown in panel a. (c) Violin plots indicating the enrichment of each gene (Man-Whitney test, −log_10_ of the P-value) in each of the indicated gene programs. Genes shown in panel b are indicated by arrows in each program. Negative control represent a randomly selected set of 200 genes.

### The stress-like program is seen at the protein level

We next sought to assess the existence of the stress-like cell state at the protein level. We performed immunofluorescence for FOS, a marker for the stress-like program (Figure 2b) on 100 human biopsy core sections (see Methods), including 62 cases of malignant melanoma, 21 metastatic malignant melanoma, and 17 nevus tissue as a control (Supplementary Figure 6).

Nuclear FOS staining was enriched in the malignant sections but not in the benign (Figure 4a). To quantify the range and intensity of FOS staining, we computed the product of the staining intensity (0-2 scale) and coverage of cancer cells (1-4 scale), yielding an index ranging from 0 to 8 (see Figure 4b for examples). Examining the distribution of the scores across the sections we found that malignant and metastatic sections have higher FOS scores compared to the benign sections (Figure 4c, *P*<10^−4^ and *P*<10^−2^, respectively). This result provides further evidence for the existence of stress-like cells in melanoma, and its enrichment in more advanced lesions (i.e., metastatic), suggesting it may mark cells contributing to poor prognosis such as invasiveness or drug resistance.

**Figure 4.**
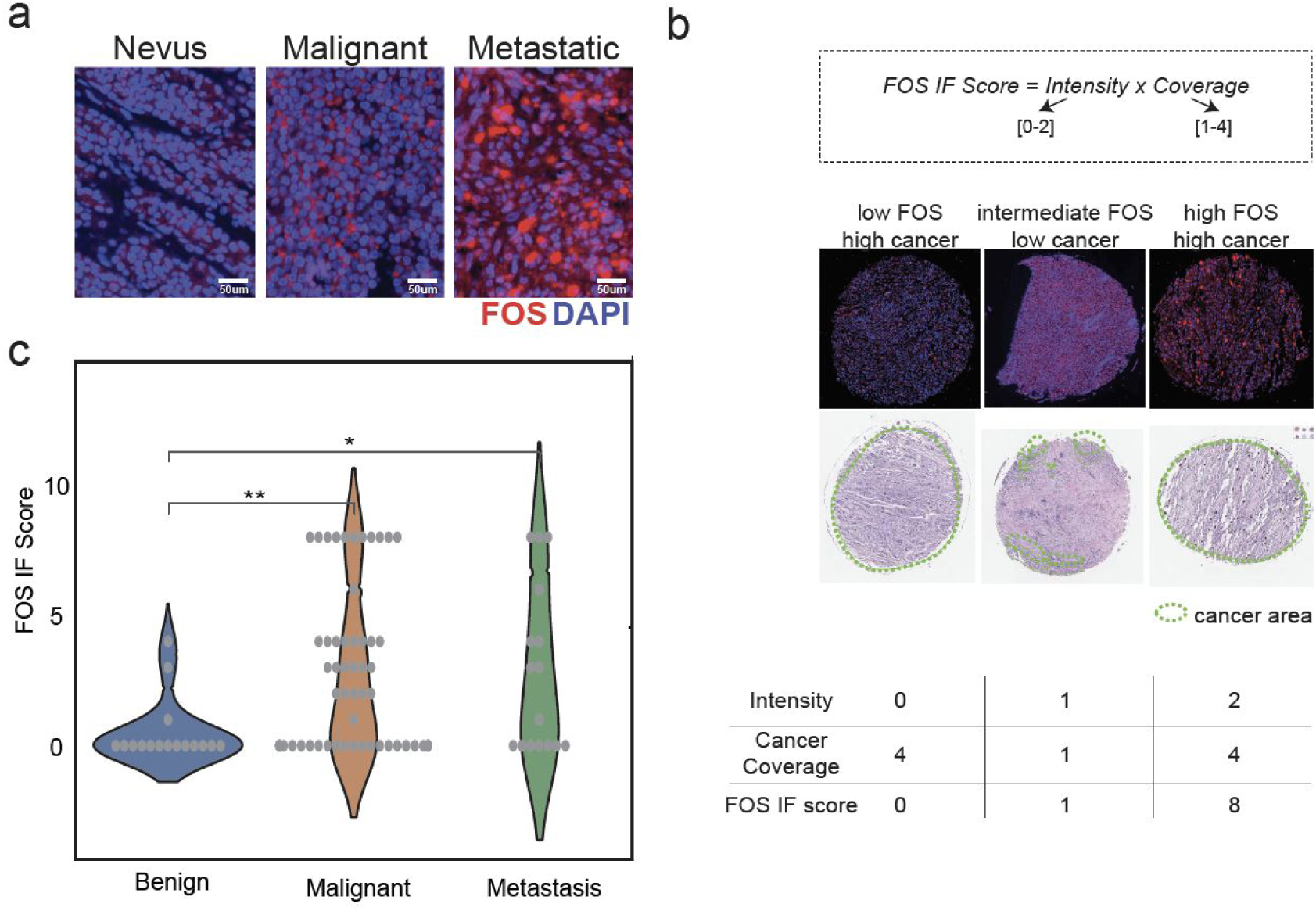
Stress-like gene program at the protein level. (a) FOS immunofluorescence images of malignant melanoma and nevus. (b) Immunofluorescence (IF) score and examples for low, medium and high FOS IF intensities and cancer coverage. (c) Violin plots of IF score distributions across 100 tumor microarray core sections. Malignant and metastasis cores have significantly high IF score compared to benign tumors (*P*<10^−4^ and *P*<10^−2^, respectively).

### The cell stress-like program is also seen in other cancer types

Since the program defined for the stress-like state (i.e., *fos*, *jun*, heat shock proteins) does not contain genes specific to melanocyte biology, in contrast to the neural crest/melanocyte states, we hypothesized that it might be conserved in other cancer types. To study this at the transcriptomics level, we applied our analytic approach to four previously published scRNA-Seq tumor datasets: triple negative breast cancer (TNBC) (Kim et al., 2018), pancreatic cancer adenocarcinoma (PDAC) (Moncada et al., 2019), oligodendroglioma (Tirosh et al., 2016b) and melanoma (Tirosh et al., 2016a). For each cancer type, we analyzed the transcriptomes of the cancer cells in isolation: 388 single cells from a TNBC patient, 462 single cells from a PDAC patient, 692 from an oligodendroglioma patient and 1257 single cells from melanoma patients, all without treatment. Studying these cancer cells using PCA, we found a triangle-shaped distribution of cells for each cancer type (Figure 5), reminiscent to that found for zebrafish melanoma (Figure 1c). Examining the expression of the stress-like gene program we found that it is enriched in one vertex in all datasets (Figure 5). In order to examine if these enrichments are statistically significant, we computed a p-value using a bootstrapping approach. We first calculated the overall expression of 10,000 randomly selected gene lists (equal in size to the stress-like program). We next assigned a p-value to each dataset by examining where the stress-like gene expression program falls within the distribution of the 10,000 lists. These confirmed that in all four datasets examined, the enrichment of the stress-like program is significant. These results provide evidence for the generality of the stress-like state across other cancers. Moreover, this suggests that the stress-like state may be conserved across diverse cancer types and may therefore play an important role in tumor progression.

**Figure 5.**
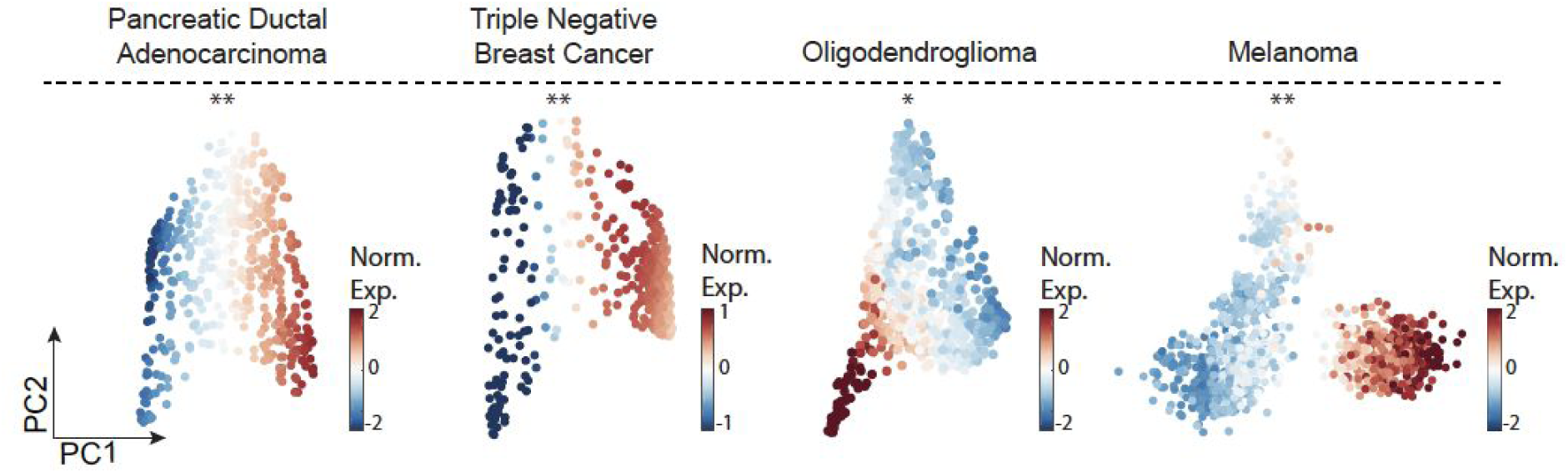
Stress-like cells are conserved across other cancer types. PCA on PDAC (Moncada et al., 2019), TNBC (Kim et al., 2018), oligodendroglioma (Tirosh et al., 2016b) and melanoma (Tirosh et al., 2016a) tumor cancer cells. Color indicates normalized expression levels of the stress-like program which is significantly enriched in one vertex (*, *P*<10^−4^; **, *P*<10^−2^).

To extend our dataset, we again used the FOS protein as a marker for stress-like cells and queried for its expression in various other cancer tissues. IF on pancreatic ductal adenocarcinoma (PDAC) showed similar patterns to melanoma where heterogeneous nuclear staining of FOS is observed across the different cancer cells (Supplementary Figure 7). Extending to other tissues, using the human protein atlas dataset (Thul et al., 2017; Uhlén et al., 2015), we noticed that heterogeneous nuclear expression of FOS is observed in other cancer types such as breast cancer, glioma, prostate cancer and head and neck cancer (Supplementary Figure 7).

### A ubb-tdTomato transgenic line as a reporter of the stress-like state

We next sought to functionally assess whether the stress-like cells in our melanomas had unique biological properties that made them particularly pro-tumorigenic. We thus built a fluorescent transgenic reporter that allowed us to isolate and characterize the stress-like cancer cells (Figure 6a). In the stress-like transcriptional program, there were three major classes of genes that we thought would be reasonable transcriptional reporters: *fosab, hsp70* or *ubb*. We attempted to make a transgenic reporter for *fos* using a promoter/enhancer fragment, but in our hands this was not a sensitive reporter of the stress state. Although *hsp70* promoter fragments have been used successfully in the fish (Venero Galanternik et al., 2016) in our hands this was found to be leaky in the melanomas and expressed in nearly all cells. We next turned to the *ubb* promoter, as a promoter fragment for this gene has been well characterized (Mosimann et al., 2011) and we found it to be a highly sensitive readout of the stress cell state, as detailed below.

**Figure 6.**
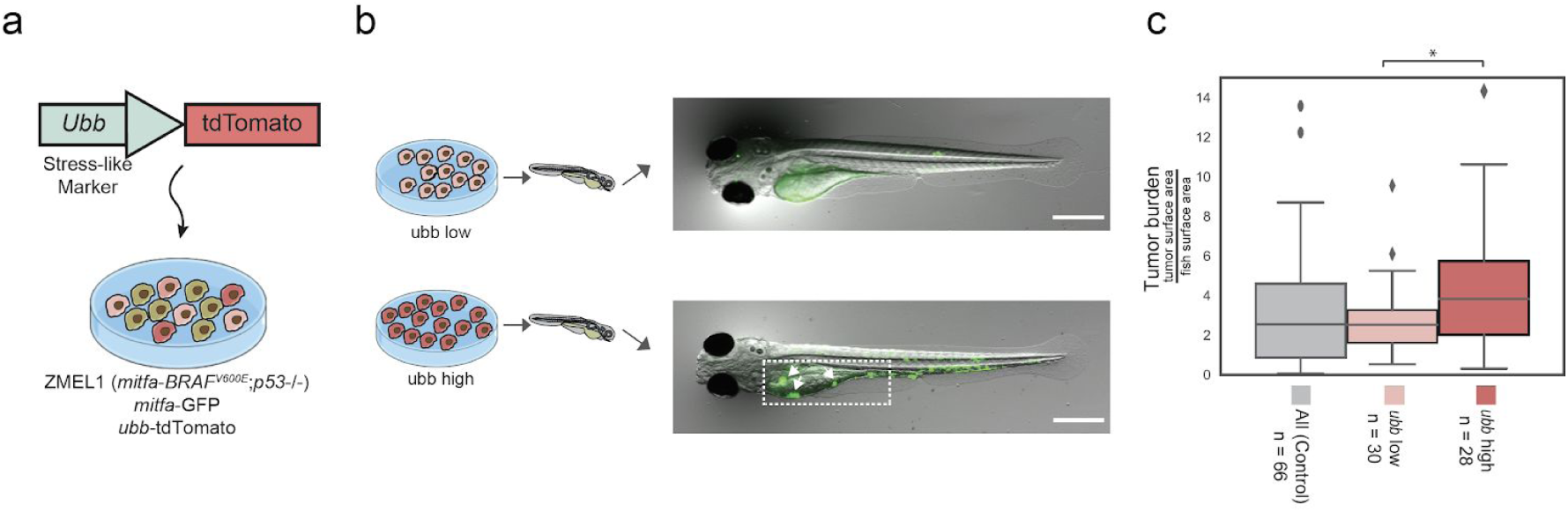
Stress-like cells form higher burden tumors. (a) The ZMEL1-GFP;*ubb*-tdTomato system to track and select for the cells in the stress-like state. (b) ZMEL1-GFP;*ubb*-tdTomato cells were sorted to high and low levels of tdTomato intensities and injected into zebrafish for tumor initiation assay followed by quantification of GFP intensity. (c) Boxplot of tumor burden quantified by GFP intensity of the two different levels of tdTomato compared to parental unsorted cells as a control (mix of population with no selection). Tumor sizes were significantly higher when high tdTomato cells were injected (Mann-Whitney test; *, *P*< 5×10^−2^).

The *ubb* gene encodes for the ubiquitin-B protein, a protein degradation system that is commonly activated under periods of diverse types of cell stress (Flick and Kaiser, 2012), including DNA damage, changes in temperature, oxidative damage, hypoxia and starvation. It serves in part to target proteins for degradation via the proteasome, including other stress genes such as *fos* that we identified in our analysis as part of the stress-like program (Stancovski et al., 1995). The ubiquitin system acts in concert with other stress related genes such as the heat shock proteins, which can act as chaperones for misfolded proteins (Dai and Sampson, 2016) and together can act as a pivotal mechanism by which the cell can either survive stress or undergo apoptosis if the stress cannot be resolved. *ubb* itself is transcriptionally induced under stress conditions such as oxidative stress, making it an ideal reporter of the stress-like state (Bianchi et al., 2015). We reasoned that cells with higher levels of *ubb* transcription would be representative of a more general stress program. We therefore hypothesized that we could use transcriptional levels of *ubb* as an indicator of the more general stress state we observed in our melanomas. To test this, we used a previously described *ubb* transcriptional promoter fragment (Mosimann et al., 2011) to drive tdTomato, and stably inserted this transgene into the ZMEL1-GFP melanoma cell described above. This *ubb* reporter then allowed us to isolate *ubb*^hi^ versus *ubb*^lo^ cells from the melanoma population. We confirmed that the transgene is only being integrated once per cell by measuring genomic *ubb* levels by qPCR. We validated that high intensity of tdTomato represents the stress-like cells in two ways: (1) Using qPCR, we found that *fosab*, *hsp70.1*, *junba* and *ubb* are more highly expressed in tdTomato-high cells compared to tdTomato-low cells, providing support for the stress-like identity (Supplementary Figure 7). We confirmed that the high fluorescence was only reflected in tdTomato and not GFP intestines to rule out autofluorescence artifacts. (2) The stress-like program includes eight heat shock genes. We induced their expression by culturing the cells in heat shock conditions (37°C overnight, compared to optimum conditions, 28°C). Examining tdTomato intensity as a measure for *ubb* expression, we detected a significant increase in heat shock conditions compared to the control (*P*<0.001, effect size = 0.8). This provides evidence for the notion that stress by heat shock leads to enrichment in *ubb* high cells (Supplementary Figure 8). From these experiments, we concluded that cells marked by high *ubb* expression are also characterized by high expression of other key stress-like program genes and that the culturing of the cells under heat shock conditions induces this state.

### The stress-like cells are more efficient at seeding new tumors

Recent work has pointed to an important role of cell stress in mediating tumor dormancy and the ability to see new tumor sites. For example, an imbalance in the ratio of ERK versus p38/stress signaling can dictate the fate choice between dormancy and proliferation (Harper et al., 2016; Ranganathan et al., 2006), and under the right microenvironment, these dormant cells can re-enter the cell cycle and begin to proliferate. Similarly, FBXW7 is a subunit of the SCF ubiquitin ligase complex that regulates HSF1 and stability of the heat shock proteins. Tumors that lose FBXW7 have elevated heat shock signatures and are more efficient at metastatic seeding (Kourtis et al., 2015), supporting the concept that stress signaling can promote tumor progression. We therefore test whether our stressed population, which occurs endogenously in the transgenic melanomas, might be more efficient at seeding new tumors. For this we developed a highly stringent assay in which we can test whether a small number of cells can give rise to a tumor after transplantation in the zebrafish. We built upon the logic of blastula transplantation, an assay that is commonly employed in developmental biology (Gansner et al., 2017) in which small numbers (5-10) of fluorescently labelled “donor” cells are transplanted into an unlabelled “recipient” animal and the fate of those cells is then tracked using in vivo imaging. We adopted this method to use with our ZMEL1-GFP;*ubb*-tdTomato cells as the donors, and transparent *casper* animals as recipients. From the ZMEL1-GFP;*ubb*-tdTomato parental population, we sorted *ubb*^hi^ versus *ubb*^lo^ cells using flow cytometry, and then transplanted 5-10 of each of these cells into recipient animals. We performed this assay with three groups of fish: 1. *ubb*^hi^ (stress-like) cells, 2. *ubb*^lo^ (non-stress-like cells), and 3. and a mix of the parental cells without prior selection. After 5 days, we then quantified the tumor burden by calculating the percentage of the entire fish that is covered by GFP+ cells intensity in a total of n=124 animals. Because the GFP is driven by the *mitfa* promoter, it is independent of *ubb* mediated transcription. This analysis revealed that overall tumor burden in animals seeded by the *ubb*^hi^ cells were larger when compared to the tumors seeded by the *ubb*^lo^ cells, or from tumors seeded by the parental unsorted cells (Figure 6c, Mann-Whitney test, *P*< 5×10^−2^). This indicates that under these highly stringent transplantation conditions, *ubb*^hi^ stress-like cells are more efficient at seeding tumors than the non-stressed isogenic melanoma cells.

### The stress-like state is associated with drug resistance

We next sought to determine whether the stress-like subpopulation of cancer cells is more drug resistant. Rambow et al. previously identified a population of persister, starved-like cells that arose after exposure to BRAF/MEK inhibitors (Rambow et al., 2018), but our data suggests that these cells may correspond to the stress-like population that is present before exposure to drug and in early stages of tumorigenesis (Figure 1e and 2d). We therefore hypothesized that the stress-like cells may be intrinsically more resistant to either BRAF or MEK inhibitors, and that these cells present in the initial tumor may give rise to the persister population. To test this, we examined the effect of BRAF or MEK inhibitors on tumor burden and populations using the ZMEL1-GFP;ubb-tdTomato cells. We transplanted 5-10 cells as above, and then treated the fish with either a BRAF inhibitor (dabrafenib) or a MEK inhibitor (trametinib). We then imaged the fish after 4 days of these treatments, and quantified the burden and the percentage of stress-like cells by measuring GFP and tdTomato intensities, respectively (Figure 7a). As a correlate to the *in vivo* imaging, we also disaggregated the entire fish at this time point and used FACS to quantify the number of *ubb*^hi^ versus ubb^lo^ cells based on tdTomato intensity. Surprisingly, we found a significant difference in response between the BRAF vs. MEK inhibitor.

**Figure 7.**
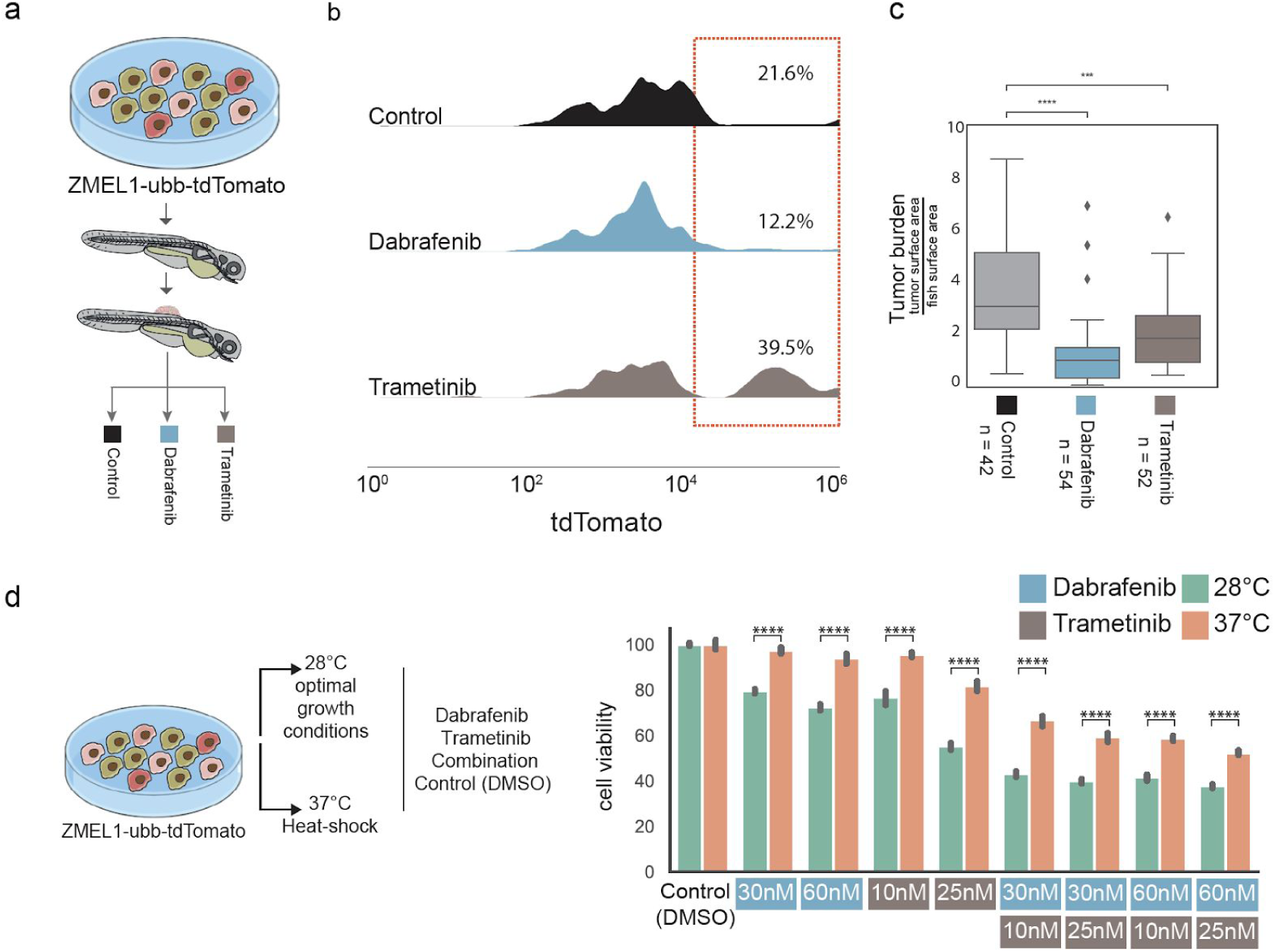
*ubb*^hi^ cells are resistant to a MEK inhibitor. (a) ZMEL1-GFP;*ubb*-tdTomato cells were injected to zebrafish embryos and the resulting tumors were treated according to the indicated conditions. (b) Histograms of tdTomato intensities, measured by FACS, of the three different treatment groups. Note the enrichment of *ubb*^hi^ cells with Trametinib-treated tumors. (c) Boxplot of tumor burden quantified by GFP intensity of the three different treatment groups. Tumor sizes were significantly smaller when treating with Dabrafenib and Trametinib (Mann-Whitney test; ****, P<10^−4^; ***, P<10^−3^, respectively). (d) Left panel, schematic of ZMEL1=GFP;*ubb*-tdTomato cells cultured in optimal and heat shock conditions and exposed to different drug concentrations. Right panel, bar plot of cell viability across culturing conditions (optimal/heat shock) and drug treatments. Cell viability is significantly higher under drug treatment when cells were cultured in heat shock conditions (Mann-Whitney test, ****, *P*<10^−4^).

We found that the BRAF inhibitor effectively suppressed both the *ubb*^hi^ and ubb^lo^ cells, suggesting the stressed versus non stressed population are equally sensitive to this targeted therapy (Figure 7b). In contrast, we found that the MEK1/2 inhibitor-treated fish effectively eliminated the ubb^lo^ cells, but failed to eradicate the *ubb*^hi^ cells (Figure 7b).

Overall, tumor burden was significantly smaller when treating with either BRAF or MEK inhibitors but the effect was largest when treated with BRAF inhibitor (Figure 7c, Mann-Whiteny test, P<5×10^−3^), pointing out the suppression of both *ubb*^hi^ and *ubb*^lo^ cells reduces tumor burden more efficiently. This observation is particularly interesting in light of recent work that shows that HSF1 is a direct phosphorylation target of MEK, and is a direct activator of the HSF1 program (Tang et al., 2015). This suggests the possibility that the stress population might have higher levels of MEK activity, which might account for their relative resistance to the MEK inhibitors. Moreover, sustained activation of ERK, which is directly downstream of MEK, leads to ubiquitin-mediated degradation of DUSP1 (also called MKP-1), which is a negative regulator of the pathway (Lin et al., 2003), providing a plausible mechanism by which the *ubb*^hi^ cells might be protected against MEK inhibition due to their higher levels of DUSP1 degradation. Why this occurs more from MEK inhibitors rather than BRAF inhibitors, which target the same pathway, remains unclear but could be due to differential feedback regulation of the MAP kinase pathway by each of these drugs, as has been previously suggested (Lito et al., 2012). These hypotheses will need further investigation in the future, but our data supports the notion that a stress-like cell state exists in the tumor even before drug exposure, and that these cells are intrinsically resistant to the MEK inhibitors commonly used in the clinic.

### Extrinsic induction of the stress-like program induces drug resistance

The above data suggest that cells with higher levels of a stress-like transcription program are intrinsically resistant to MEK inhibitors. We next queried whether extrinsic induction of the stress-like state is sufficient to trigger this resistance. As noted above, the heat shock program is a direct target of MEK, and given our finding above that heat shock of the melanoma cells was sufficient to trigger increased expression of the ubb reporter (Supplementary Figure. 8), we reasoned that heat shock of the cells might affect sensitivity to BRAF or MEK inhibitors. To test this, we grew ZMEL1-GFP;*ubb*-tdTomato cells at either 28°C (the normal zebrafish temperature) or at 37°C (a temperature long known to induce the heat shock response in zebrafish, Figure 7d, left panel). Strikingly, this induced resistance to both BRAF as well as MEK inhibitors across a wide range of doses and combinations (Figure 7d, right panel). To further confirm that this was not an effect specific to the zebrafish, we repeated this experiment using human A375 melanoma cells, which harbor the same *BRAF^V600E^* mutation as the ZMEL1 cells. In this case, we grew the human cells either at 37°C (normal human body temperature) or at 42°C (known to induce heat shock in human cells) and found a similar phenomenon - cells at the higher temperature were more resistant to both BRAF or MEK inhibitors across the doses tested (Supplementary Figure 9). Collectively, these data further support a model in which a stress-like state, whether intrinsic to the cell or extrinsically induced, is associated with resistance to these MAP kinase inhibitors.

## DISCUSSION

Here we have studied the cells states of cancer cells in zebrafish and human melanomas. We detected three recurring gene expression programs underlying these states: a mature melanocyte, a neural-crest and a stress-like transcriptional program. Other recent findings using scRNA-seq have also detected transcriptional programs in melanoma cancer cells. Tirosh et al. showed that cancer cells have distinct transcriptional programs that capture the known proliferative and invasive states (Tirosh et al., 2016a), Rambow et al. identified four transcriptional programs in patient-derived xenografts (PDX) formed by BRAF mutant patients that were treated with RAF/MEK inhibition drugs (Rambow et al., 2018) and Tsoi et al. found that bulk melanoma tumors show different states along the trajectory of differentiation (Tsoi et al., 2018). While the other studies have also noted the stress-like state, they did not pursue it functionally, and a concern in the field has been that scRNA-Seq data may artificially exhibit this state because of cell dissociation and sorting methods (van den Brink et al., 2017). Our focused analysis on the stress-like cell state led us to conclude that it is a bona fide component of the tumor and that it is has tumor-promoting and drug tolerant properties (Figures 6 and 7). While our studies support the existence of this population in our samples, it is likely that the artifactual observations of stress states previously shown in single cell data is relevant depending upon the particular samples and preparation methods. We cannot exclude the possibility that in some studies, observation of a stress-like state is indeed an experimental artifact, but these will need to be evaluated on a case by case basis. In this Discussion, we consider the general occurrence of the stress-like state across cancer types, its possible role, qualities and implications for treatment.

As evidence that the stress-like state is part of the tumor - beyond the single cell transcriptomics in our zebrafish melanoma and those of Rambow et al and Tirosh et al. - we report data from spatial transcriptomics and immunofluorescence. Both of these technologies have the advantage that they do not rely on dissociation of the tissue which may lead to artifactual induction of a stress-like state. Indeed, in our analysis of 100 sections of various stages of melanoma, we find that FOS protein expression is significantly enriched in malignant cells versus benign lesions. Further evidence of its comes from our observations that the stress-like program is conserved across both species (Figure 2) and distinct cancer types (Figure 5), which we examined using human PDAC sections, human protein atlas and published scRNA-seq datasets (Figure 4).

What role does the stress-like state play in the tumor? Cell stress is increasingly linked to phenotypes such as metastasis and drug resistance, but the term stress can refer to a wide array of both inducers and mediators. In cancer, malignant phenotypes are linked to processes such as DNA damage, reactive oxygen species, endoplasmic reticulum (ER) stress, and starvation stress among others. What is common to many of these different forms of stress is that they often cause induction of a set of downstream genes that allow the cell to cope with these insults. Amongst those common mediators are many of the genes we identified in our studies, and they have mechanistic inter-relationships. The *fos/jun* family are classical “immediate early response” genes that can be rapidly transcribed and act as transcription factors (Healy et al., 2013). The *fos/jun* family are regulated by both the heat shock and ubiquitin proteins, two other families we identified in our gene sets. For example, *c-fos* is degraded by ubiquitination (Stancovski et al., 1995), but at the same time, induction of heat shock proteins has been demonstrated to increase DNA binding of these factors (Mattson et al., 2004). *jun* itself is known to be a direct target of the heat shock *HSF1* factor (Sawai et al., 2013), so it is perhaps not surprising that in our findings we see elevation of both heat shock and *jun* in this cell state.

One important clinical implication of this study is that the existence of the stress program can have consequences for the intrinsic responsiveness to therapy. In melanoma, for example, HDCA8 has been shown to mediate drug resistance under multiple stress conditions such as BRAF-MEK treatment, hypoxia, UV and radiation through regulation of MAPK, decrease in *jun* acetylation and enrichment of AP-1 signaling (Emmons et al., 2019). HSF1 has been demonstrated to be essential for chemotherapeutic agent-induced cytoprotective autophagy by directly binding to ATG7 in breast cancer (Desai et al., 2013) and active in colon and lung tumors as well (Mendillo et al., 2012). Our study also helps explain prior work in which it has been observed that high levels of the heat shock protein HSP90 can promote BRAF oncogenic activity, since it can bind to and stabilize the mutant BRAF^V600E^ protein (Grbovic et al., 2006). The increased resistance to both BRAF and MEK inhibitors upon application of heat shock may be due to this stabilization of BRAF and increased MAP kinase signaling. Whether other physiologic conditions associated with increased heat, i.e. fever that occurs in cancer patients, affects response to these therapies remains an open question. In addition to drug responsiveness, it is also likely that the stress program could play a role in metastasis. Treatment with BRAF inhibitors induces a stress like phenotype with increased RhoA activity (Klein and Higgins, 2011), a gene highly associated with metastatic capacity. Increased oxidative stress is associated with invasiveness (Taddei et al., 2012) and higher levels of Wnt5A are associated with a slow cycling, nearly senescent type of cell that has invasive characteristics (Webster et al., 2015). Recent work has highlighted that translational reprogramming induces a “pseudo-starvation” state, which may be mediated by the ER stress response, as a central element of invasion (Falletta et al., 2017; García-Jiménez and Goding, 2019). Whether the stress population we identified in our studies has increased metastatic capacity remains to be determined.

One of our main findings is that the stress-like state is a consistent component of the cancer cell population, detectable at very early stages of tumorigenesis. This raises the question of what adaptive advantages this program might have for tumor initiation. The *fos/jun* pathway is a critical downstream mediator of MAP kinase signaling (Dunn et al., 2005; O’Donnell et al., 2012), suggesting that activation of BRAF^V600E^ itself may be inducing this state. However, it is likely that other factors such as hypoxia are playing a role as well, during these early stages of tumorigenesis (Webster et al., 1993). In conditions of increasing stress, in which there may be highly elevated levels of abnormally folded proteins (Santagata et al., 2012; Whitesell and Lindquist, 2005; Whitesell et al., 2014) both the heat shock and ubiquitin systems must be coordinated, the former to allow for chaperoning of those proteins and the latter for degradation of these abnormally proteins. The stress population in cancer is likely enacting a set of regulatory mechanisms that balance cell survival/quiescence versus apoptosis, which is critical in the early stages of tumorigenesis. The fact that we see the emergence of this population early in tumorigenesis, even in the absence of drug or other selection factors leads us to speculate that these states could be generated by stable epigenetic mechanisms, rather than genetic mechanisms, since we see it in multiple cancers with very different DNA mutational events (i.e. BRAF, KRAS, CDKN2A). The specific factors that induce the stress state, whether cell intrinsic or microenvironmental, will need further analysis in the future, since they may offer an opportunity for eliminating these intrinsically drug resistant “seeds” even before therapies are applied.

## DATA AVAILABILITY

The complete data that support the findings of this study have been deposited in NCBI GEO database with the accession code GSE115140.

## ACKNOWLEDGMENTS

We thank Naftalie Senderovich, and Anna Yeaton for work on the initial pilot of the project. We thank Matt Maurano and Megan Hogan for assistance with sequencing. We also thank the NYU Langone Genome Technology Center for assisting with the whole genome sequencing.

## AUTHOR CONTRIBUTIONS

I.Y. and R.M.W. conceived the project. M.B. led the collection and sequencing of the single-cell RNA-Seq data collection, with contributions from I.S.K. and N.R.C.. R.M. and M.H. contributed the spatial transcriptomics data and Y.Y contributed bioinformatics. M.T, M.H and M.B performed the functional characterization. M.B. led the analysis of the data, with significant contribution from I.Y.. I.Y. and R.M.W. provided project coordination. I.Y., R.M.W and M.B. drafted the manuscript on which all authors commented.

---

**Table S1.**
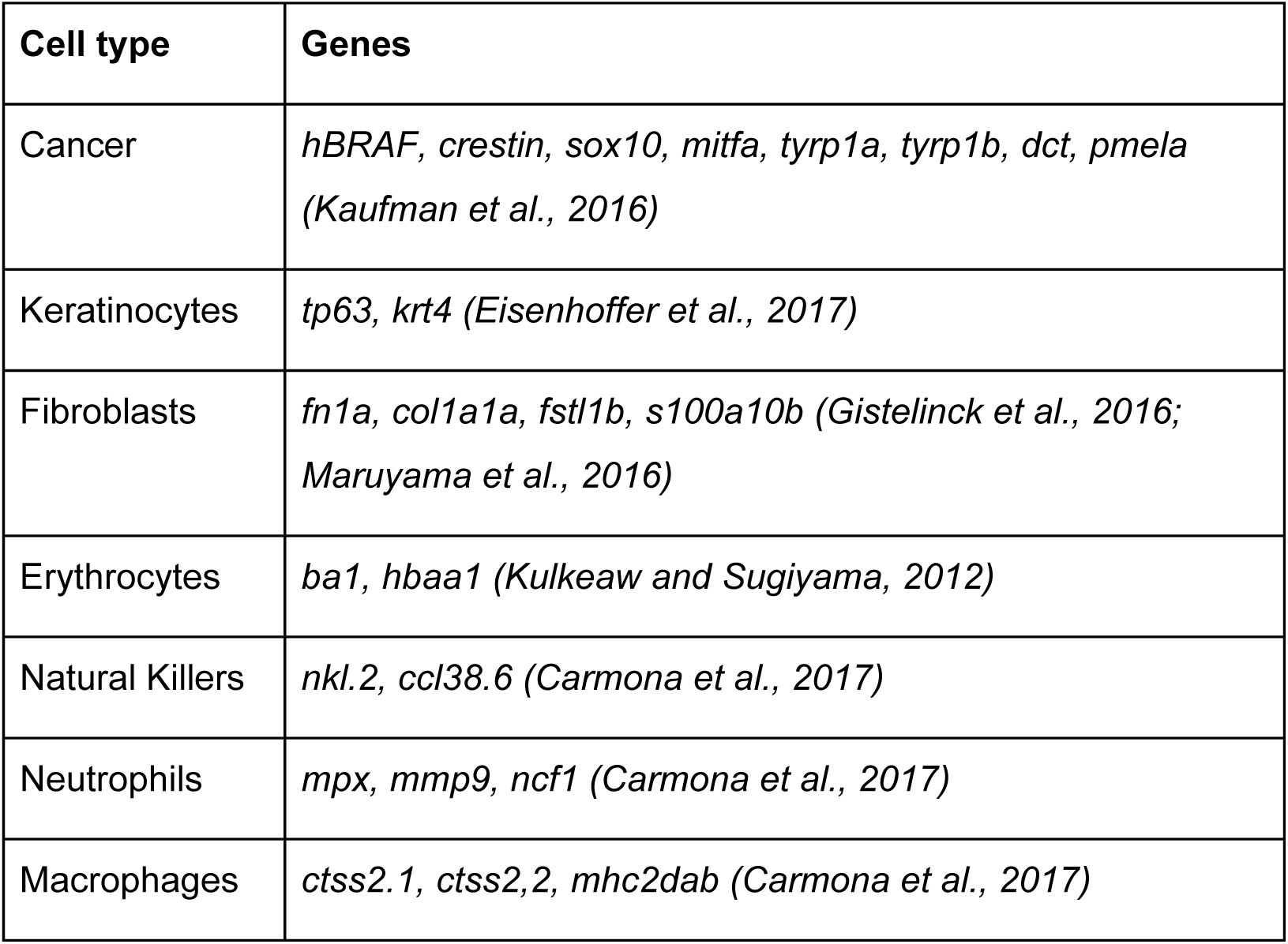
Marker genes used in analysis are indicated for each cell type.

**Table S2.**
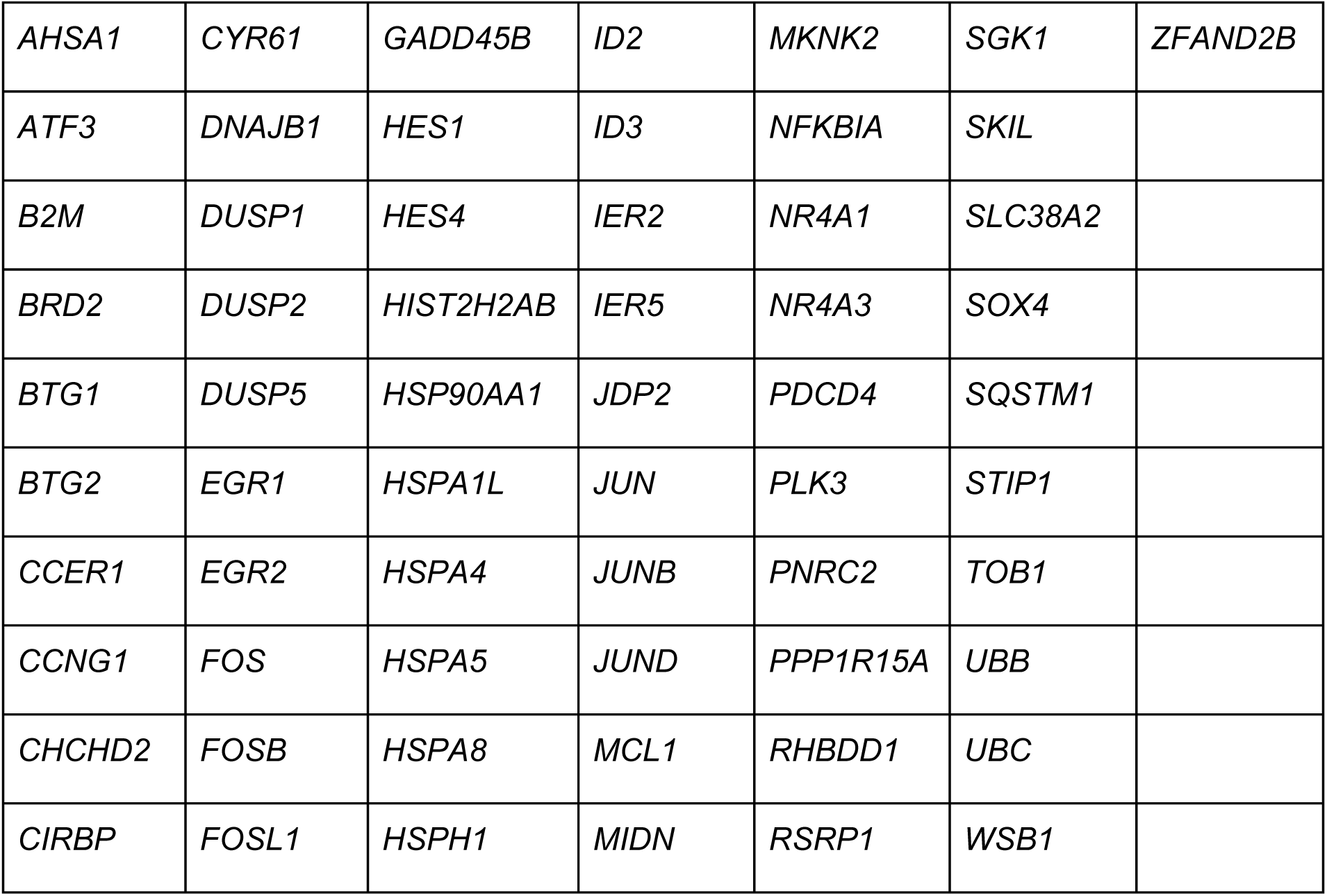
Stress-like program genes.

|  |  |  |  |  |  |  |
| --- | --- | --- | --- | --- | --- | --- |
| *AHSA1* | *CYR61* | *GADD45B* | *ID2* | *MKNK2* | *SGK1* | *ZFAND2B* |
| *ATF3* | *DNAJB1* | *HES1* | *ID3* | *NFKBIA* | *SKIL* |  |
| *B2M* | *DUSP1* | *HES4* | *IER2* | *NR4A1* | *SLC38A2* |  |
| *BRD2* | *DUSP2* | *HIST2H2AB* | *IER5* | *NR4A3* | *SOX4* |  |
| *BTG1* | *DUSP5* | *HSP90AA1* | *JDP2* | *PDCD4* | *SQSTM1* |  |
| *BTG2* | *EGR1* | *HSPA1L* | *JUN* | *PLK3* | *STIP1* |  |
| *CCER1* | *EGR2* | *HSPA4* | *JUNB* | *PNRC2* | *TOB1* |  |
| *CCNG1* | *FOS* | *HSPA5* | *JUND* | *PPP1R15A* | *UBB* |  |
| *CHCHD2* | *FOSB* | *HSPA8* | *MCL1* | *RHBDD1* | *UBC* |  |
| *CIRBP* | *FOSL1* | *HSPH1* | *MIDN* | *RSRP1* | *WSB1* |  |

## METHODS

### Fish strains and handling

*minicoopr* fish were generated as previously described (Ceol et al., 2011; Iyengar et al., 2012). Fish with the genotype *mitfa-BRAF^V600E^*;*p53*-/-;*mitfa*-/- were incrossed. 1-cell stage embryos were injected with a plasmid containing *mitfa-MITF* and *mitfa-GFP*. Fish were screened at 3 days for melanocyte rescue, visualized as black spots along the skin. The fish were raised to adulthood (4-12 months) and screened for the appearance of GFP-positive tumors. For biopsy experiments, fish with visible tumors were anesthetized with Tricaine (MS222), and a biopsy was taken using a 1mm biopsy punch. The fish were then allowed to recover and were returned to the main aquatics system.

### Single-cell RNA-seq collection and processing

#### Tumor biopsy sample collection

Biopsy samples were obtained from the same location of the tumor and placed in a 1.5mL Eppendorf tube followed by the addition of 500µL 0.25% Trypsin-EDTA for digestion. The digestion was carried out at 37°C in a thermomixer for 15-30 min to soften the tissue, with tissue mashings every 5 minutes using a disposable pestle to break up softened tissue. Upon completion of incubation at 37°C, 500µL of DMEM10 were added to deactivate the trypsin. Cells were washed three times by spinning down the sample at 500 rcf for 5 minutes and resuspended in PBS. The sample was then filtered twice using 5mL polystyrene round-bottom tube with 35µm cell-strainer. Viability and single cell consistency were checked prior to encapsulation of the cells with the inDrop system (Klein et al., 2015; Zilionis et al., 2017) for each biopsy taken.

#### Single-cell encapsulation, processing and bioinformatics pipeline

inDrop encapsulation of the cells and reverse transcription (RT) reaction was carried out as previously described (Klein et al., 2015; Zilionis et al., 2017). RNA amplification and library preparation was carried out according to this protocol incorporating the changes introduced in Zilionis et al. 2017 based on the CEL-Seq2 protocol (Hashimshony et al., 2016) on batches of 1,500-3,000 cells from each biopsy taken. The number of PCR cycles required for final library amplification ranged from 9-13 cycles. Single-cell RNA-Seq library sequencing was carried out using the Illumina NextSeq 500/550 machine. Pair-end sequencing was carried out with read1 (barcodes) for 34bp, index read for 6bp and read2 (transcripts) for 50bp. Raw sequencing data obtained from the inDrop method was processed using a custom-built pipeline, (available at https://github.com/flo-compbio/singlecell). The location of the known “W1” adapter sequence of the inDrop RT primer, was located in the barcode read (read 2). Reads for which the W1 sequence could not be detected were discarded. The start position of the W1 sequence was then used to infer the length of the first part of the inDrop cell barcode in each read, which can range from 8-11bp, as well as the start position of the second part of the inDrop cell barcode, which is 8bp long. Cell barcode sequences were mapped to the known list of 384 barcode sequences for each read. The resulting barcode combination was used to identify the cell from which the fragment originated. Finally, UMI sequence was extracted, and reads with low confidence base calls for the six bases comprising the UMI sequence (minimum PHRED score less than 20) were discarded. The reads containing the mRNA sequence (read 1) were mapped using STAR with parameter “—outSAMmultNmax 1” and default settings otherwise (Dobin et al., 2013). Expression was quantified by counting the number of reads mapped to each gene. The genome and gff file used included the zebrafish genome (Version 10 (Zerbino et al., 2018)) and the BRAF human vector.

#### Quality control and filtering of low-quality cells

Single-cell transcriptomes with UMIs>750, mitochondrial transcripts < 20% and ribosomal transcripts < 30% were retained for analysis, leaving 7,278 cells. The same approach was applied for the other tumors leaving 1171 and 1563 cells, respectively, yielding a total of 10,012 (out of 15,000 processed for all samples). Expression profiles were smoothed using MAGIC (van Dijk et al., 2017) with parameter k = 7 to reduce noise after optimization using different k values. UMI counts were normalized by the total number of transcripts per cell, and a scale factor equivalent to the median number of transcripts for that cell was applied (transcripts per median, TPM). Expression was transformed using Freeman-Tukey transform (FTT) as described previously (Wagner et al., 2017).

### Cell type clustering

Clustering was performed by first distinguishing the cancer cells from non-cancer cells by detecting the expression of the human BRAF gene. Next, hierarchical clustering was performed on the non-cancer cells with Ward’s criterion using the most variable genes (defined as Fano factor and mean expression above mean-dependent thresholds). Clustering was performed over correlations computed from the smoothed expression of the selected genes (Z-score of the TPM). This process initially revealed five clusters: cancer, immune, keratinocytes, fibroblasts and erythrocytes. After examining the immune cluster using recently published markers (Carmona et al., 2017) we could separate the immune cluster into three sub-clusters: macrophages, natural killers and neutrophils. To identify each cluster, we obtained a list of marker genes by examining genes that are differentially expressed (P<10^−6^, Kolmogorov-Smirnov test; effect size >0.2, Cohen’s d).

### Dimensionality Reduction (PCA and tSNE)

Dimensionality reduction methods were performed on TPM transformed data using variable genes (defined as Fano factor and mean expression above mean-dependent thresholds). tSNE was performed using the following parameters: perplexity = 30 and initial dimension = number of principal components explaining >90% of the variance (Maaten and Hinton, 2008).

### Cancer cell type analysis

To find genes that are uniquely expressed in each cancer cell type we first identified each vertex: 1. low pc2 score, 2. high pc2 score, and 3. high pc1 score. Then we identified the 500 closest cells (Euclidean distance) to create 3 groups of cells. For each gene we then checked if its expression is significantly higher in one group compared to the other two and higher in that program (P < 10^−10^, Kolmogorov-Smirnov test, effect size > 0.2, Cohen’s d). As a control we also used a different approach which yielded similar results. This included first identifying dynamic genes (defined as Fano factor and mean expression above mean-dependent thresholds) followed by unsupervised clustering to identify 3 clusters. We found a significant overlap between the set of genes differentially expressed genes in these clusters with the ones we had originally identified using the approach described first.

### Re-analysis of Tirosh et al. human scRNA-Seq melanoma dataset

Normalized scRNA-Seq data was retrieved from the Tirosh et al. publication (Tirosh et al., 2016a) and transformed using the Freeman-Tukey approach (Wagner et al., 2017). We examined the cancer cells as annotated by Tirosh et al., and of those only the MITF high (proliferating) cells.

### Re-analysis of Mica et al. human melanocyte differentiation dataset

Normalized microarray data was retrieved from Mica et al. (Mica et al., 2013) for the wild-type cell line studied using the standard and neural crest optimized protocols. PCA was performed on samples using the most variable genes (defined above). K-means clustering with K=3 yielded the following groupings: samples from day 0 to day 3, samples from day 6 to day 11 and primary and mature melanocyte. To calculate the Pearson correlation of each cancer cell state to each grouping of melanocyte differentiation, in silico bulk expression profiles were created by averaging each cancer cell type from the human scRNA-Seq dataset (Tirosh et al., 2016a). Next, the resulting profiles were scaled to the same range of melanocyte differentiation data set using min/max scaling. Finally, only genes that are differentially expressed among the three groupings were used for the correlation calculation (P < 10^−5^, Kolmogorov-Smirnov test and expression above a mean dependent threshold).

### Re-analysis of Moncada et al. human scRNA-Seq pancreatic adenocarcinoma dataset

scRNA-Seq data was retrieved from Moncada et al. publication (Moncada et al., 2019), smoothed using the MAGIC method (van Dijk et al., 2017) and transformed using the Freeman-Tukey approach (Wagner et al., 2017). We examined only the of cancer cells, 462 in total, using the approach described in “Dimensionality Reduction (PCA and tSNE)” part. The stress program was defined using only the genes that are enriched for this cancer cell type (see “Cancer cell type analysis” part for details) in the human single-cell RNA-Seq Melanoma dataset.

### Re-analysis of Kim et al. human scRNA-Seq triple negative breast cancer dataset

Normalized scRNA-Seq data was retrieved from Kim et al. publication (Kim et al., 2018), smoothed using the MAGIC method (van Dijk et al., 2017) and transformed using the Freeman-Tukey approach (Wagner et al., 2017). We examined only the of cancer cells from donor P_6 before treatment, using the same approach as described in “Re-analysis of Moncada et al. human scRNA-Seq pancreatic adenocarcinoma dataset”.

### Re-analysis of Tirosh et al. human scRNA-Seq oligodendroglioma dataset

Normalized scRNA-Seq data was retrieved from Tirosh et al. publication (Tirosh et al., 2016b), smoothed using the MAGIC method (van Dijk et al., 2017) and transformed using the Freeman-Tukey approach (Wagner et al., 2017) The stress-like transcriptional program was identified using the same approach as described in “Re-analysis of Moncada et al. human scRNA-Seq pancreatic adenocarcinoma dataset”.

### Immunofluorescence of FOS protein on tumor microarray

Tumor microarrays (TMA) were obtained from US Biomax (ME1004g). The slide was baked for 30 minutes in 60°C and washed three times with Xylene, 100% EtOH and 95% EtOH. After rinsing with DI H20, antigen retrieval was performed for 12 minutes in boiling TE buffer. Slide was cooled down and rinsed with DI H20 following a wash in TBS+0.05% Tween. The array was stained for FOS (1:1000; Synaptic systems 226 003) for 48 hours at room temperature in a wet chamber. Before applying the secondary antibody, three washes with TBS+0.05% Tween were performed. DAPI and Secondary antibody (1:100) was applied and incubated for 1hr in a wet chamber at room temperature followed by 3 washes in TBS+0.05% Tween. Slide was dried and mounted before scanning for imaging.

### Spatial transcriptomics (ST) of zebrafish tumors

#### Tissue preparation, cryosectioning, fixation, staining, and brightfield imaging

Zebrafish melanoma tumors were obtained by sectioning the entire tumor with its surrounding tissue. Tissue was transferred from 1X-PBS to a dry, sterile 10-cm dish and gently dried prior to equilibration in cold OCT for 2 minutes. The tissue was then transferred to a tissue-mold with OCT and snap-frozen in liquid nitrogen-chilled isopentane. Tissue blocks were stored at −80°C until further use. Prior to cryosectioning, the cryostat was cleaned with 100% ethanol, and equilibrated to an internal temperature of −18°C for 30 minutes. Once equilibrated, OCT embedded tissue blocks were mounted onto the chuck and equilibrated to the cryostat temperature for 15-20 minutes prior to trimming. ST slide was also placed inside cryostat to keep the slide cold and minimize RNase activity. Sections were cut at 10 µm sections, mounted onto the ST arrays, and stored at −80°C until use, maximum of two weeks. Prior to fixation and staining, the ST array was removed from the −80°C and into a RNase free biosafety hood for 5 minutes to bring to room temperature, followed by warming on a 37°C heat block for 1 minute. Tissue was fixed for 10 minutes with 3.6% formaldehyde in 1X PBS, and subsequently rinsed in 1x PBS. Next, the tissue was dehydrated with isopropanol for 1 minute followed by staining with hematoxylin and eosin. Slides were mounted in 65 µl 80% glycerol and brightfield images were taken on a Leica SCN400 F whole-slide scanner at 40X resolution.

#### Spatial Transcriptomics (ST) barcoded microarray slide information

Library preparation slides used were purchased from Spatial Transcriptomics (https://www.spatialtranscriptomics.com; lot 10002). Each of the spots printed onto the array is 100 µm in diameter and 200 µm from the center-to-center, covering an area of 6.2 by 6.6 mm. Spots are printed with approximately 2 × 10^8^ oligonucleotides containing an 18-mer spatial barcode, a randomized 7-mer UMI, and a poly-20TVN transcript capture region (Ståhl et al., 2016).

#### On-slide tissue permeabilization, cDNA synthesis, probe release

After brightfield imaging, the ST slide was prewarmed to 42°C and attached to a pre-warmed microarray slide module to form reaction chambers for each tissue section. The sections were pre-permeabilized with 0.2 mg/ml BSA and 200 units of collagenase diluted in 1X HBSS buffer for 20 minutes at 37°C and washed with 100 µl 0.1X SSC buffer twice. Tissue was permeabilized with 0.1% pepsin in HCl for 4 minutes at 42°C and washed with 100 µl 0.1X SSC buffer twice. Reverse transcription (RT) was carried overnight (∼18-20h) at 42°C by incubating permeabilized tissue with 75 µl cDNA synthesis mix containing 1X First strand buffer (Invitrogen), 5 mM DTT, 0.5 mM each dNTP, 0.2 µg/µl BSA, 50 ng/µl Actinomycin D, 1% DMSO, 20 U/µl Superscript III (Invitrogen) and 2U/µl RNaseOUT (Invitrogen). Prior to removal of probes, tissue was digested away from the slide by incubating the tissue with 1% 2-mercaptoethanol in RLT buffer (Qiagen) for one hour at 56°C with interval shaking. Tissue was rinsed gently with 100 µl 1X SSC, and further digested with proteinase K (Qiagen) diluted 1:8 in PKD buffer (Qiagen) at 56°C for 1 hour with interval shaking. Slides were rinsed in 2X SSC with 0.1% SDS, then 0.2X SSC, and finally in 0.1X SSC. Probes were released from the slide by incubating arrays with 65 µl cleavage mix (8.75 μM of each dNTP, 0.2 μg/μl BSA, 0.1 U/μl USER enzyme (New England Biolabs) and incubated at 37 °C for 2 hours with interval mixing. After incubation, 65 µl of cleaved probes was transferred to 0.2 ml low binding tubes and kept on ice.

#### ST library preparation and sequencing

Libraries were prepared from cleaved probes as previously described, with the following changes. After RNA amplification by in vitro transcription (IVT) and subsequent bead clean-up, second RT reaction was performed using random hexamers, eliminating the need for a primer ligation step as described previously (Hashimshony et al., 2016).

#### ST spot selection and image alignment

Upon removal of probes from ST slide, the slide is kept at 4°C for up to 3 days. The slide was placed into a microarray cassette and incubated with 70 µl of hybridization solution (0.2 µM Cy3-A-probe, 0.2 µM Cy3 Frame probe, in 1X PBS) for 10 minutes at room temperature. The slide was subsequently rinsed in 2X SSC with 0.1 % SDS for 10 minutes at 50°C, followed by one-minute room temperature washes with 0.2X SSC and 0.1X SSC. Fluorescent images were taken on a Hamamatsu NanoZoomer whole-slide fluorescence scanner. Brightfield images of the tissue and fluorescent images were manually aligned with Adobe Photoshop CS6 to identify the array spots beneath the tissue.

#### ST library sequence alignment and annotation

The raw paired end sequencing file was processed by custom pipeline CEL-Seq2 (https://github.com/yanailab/celseq2) to generate the UMI-count matrix for 1007 spots. In general, the CEL-Seq2 was adapted to spatial-transcriptomics data in 3 steps: 1) Tagging and demultiplexing. The leftmost 25nt of R1 sequence consist of 18nt for spot-specific barcode and then 7nt for UMI. R2 sequence reads contains the transcript information and its leftmost 35nt were used for mapping. The name of every R2 read is tagged with spot-specific barcode and UMI sequence that are extracted from the paired R1 read. R2 reads are demultiplexed to create the 1007 spot-specific FASTQ files. If the detected spot-specific barcode of a read is not present in the pre-defined barcodes list, the read is excluded from the downstream analysis. 2) Alignment of demultiplexed FASTQ files using Bowtie2 version 2.3.1 (Langmead and Salzberg, 2012). 3) Counting UMI using customized HTSeq (Anders et al., 2014). The reads that are aligned to are collapsed to count only once if they have the same UMI.

#### Analysis of ST data

UMI counts in each spot were normalized by the total number of transcripts per spot and then multiplied by a scale factor equivalent to the median number of transcripts per spot (TPM). A pseudocount of 1 was added prior to log10 transformation. To distinguish between cancer and non cancer areas clustering was performed as described previously (Moncada et al., 2019) and enrichment of each gene in each of the transcription programs was calculated by Wilcoxon rank sum test.

### Stress-like cells in vivo and in vitro functional assays

#### Cell lines

A375 (human melanoma cell line) was obtained from ATCC. ZMEL1 cells were generated as previously described (Heilmann et al., 2015).

#### ZMEL1-tdtomato-ubb cell line line derivation

For the generation of ZMEL1 cells expressing tdTomato under the control of the ubb promoter, LR gateway cloning was performed with 5’ubb promoter, middle entry tdTomato-NTR and 3’ SV40 fragment. Following successful cloning, 8 million ZMEL1 cells were electroporated with 15 µg of the plasmid using the Neon electroporator. Following electroporation, cells were allowed to recover for 72hr and subsequently grown in 4µg/ml blasticidin containing media for 3 weeks to select for stable integration of the plasmid.

#### Heat shock and drug treatments

ZMEL1 cells were grown at 28°C (optimum temperature) or 37°C (heat shock) for 48hr with 30, 60, 600 and 6000nM of dabrafenib or 10, 25 and 50nM of trametinib, either alone or in combination. For human melanoma cell line A375, cells were grown either at 37°C (optimum temperature) or 42°C (heat shock) and treated with the same drug concentrations as above. Cell viability was measured using the Promega CellTiter-Glo®2.0 Cell Viability Assay and luminescence was measured using a Biotek plate reader.

#### Zebrafish blastula transplants and microscopy

ZMEL1 cells expressing mitf-GFP and ubb-tdTomato were grown as previously described (Heilmann et al., 2015). Cells were trypsinized and resuspended in Dulbecco’s 1X PBS to a concentration of 2×10^7^cells/mL. Approximately 20 cells in 1nL PBS were injected into the blastula of pre-epiboly casper embryos (∼2.5-4 hours post-fertilization) using a quartz microneedle. Embryos were grown in E3 for 24 hours before adding drugs. All fish were grown at 28.5°C for the duration of the experiment. For microscopy, fish were anesthetized in Tricaine and placed on a petri dish containing 2% agarose. The fish were imaged using a Zeiss AxioZoom V16 fluorescence stereoscope with a 0.6X lens. Each fish was consecutively imaged with brightfield, GFP and Rhodamine filters. Raw images (CZIs) of each larva were exported for downstream analysis in MATLAB.

#### Drug treatments

Compounds used were dabrafenib (working concentration: 1µM; Selleckchem #S2807) and trametinib (working concentration: 25nM; Selleckchem #2673). DMSO in E3 was used as the vehicle for all drugs and for all drug controls. Drugs were added at 24 hours post-fertilization and changed every subsequent 24 hours until imaging on day 5.

#### Quantification of tumor burden

Quantifications of tumor area were performed using the MATLAB Image Processing Toolbox and a fully automated custom image analysis pipeline. Brightfield images were automatically segmented to calculate the area of the entire larva. The brightfield segmentation of the whole larva was then applied as a mask to the corresponding images of GFP fluorescence from the same fish to crop the images and reduce background noise. To measure tumor area, tumor images were thresholded and the number of pixels above the threshold within the body of the larva were counted. Images of GFP fluorescence were used to calculate tumor area for all groups, due to consistency in mitf-GFP fluorescence in all tumor cells across all groups. Tumor surface area was normalized to the total fish surface area.

#### Quantification of ubb cells using flow cytometry

For each cell group of treatment, all larval fish were disaggregated in trypsin-EDTA by shaking at 37°C for 15 minutes with intermittent gentle agitation with a Disposable Pellet Pestle (Fisher Scientific) every 5 minutes. Cells were pelleted by centrifugation and resuspended in DMEM supplemented with 2% FBS. All samples were filtered through 40μm cell strainers to achieve single cell suspensions prior to FACS. Individual cancer cells, from each treatment group, were selected based on GFP expression using FACS (Fluorescence activated cell sorting) SONY SH800 cell sorter. The distribution of tdTomato was recorded and analyzed using FlowJo software v10.6.0 (Tree star, Ashland, OR, USA).

**Supplementary Figure 1.**
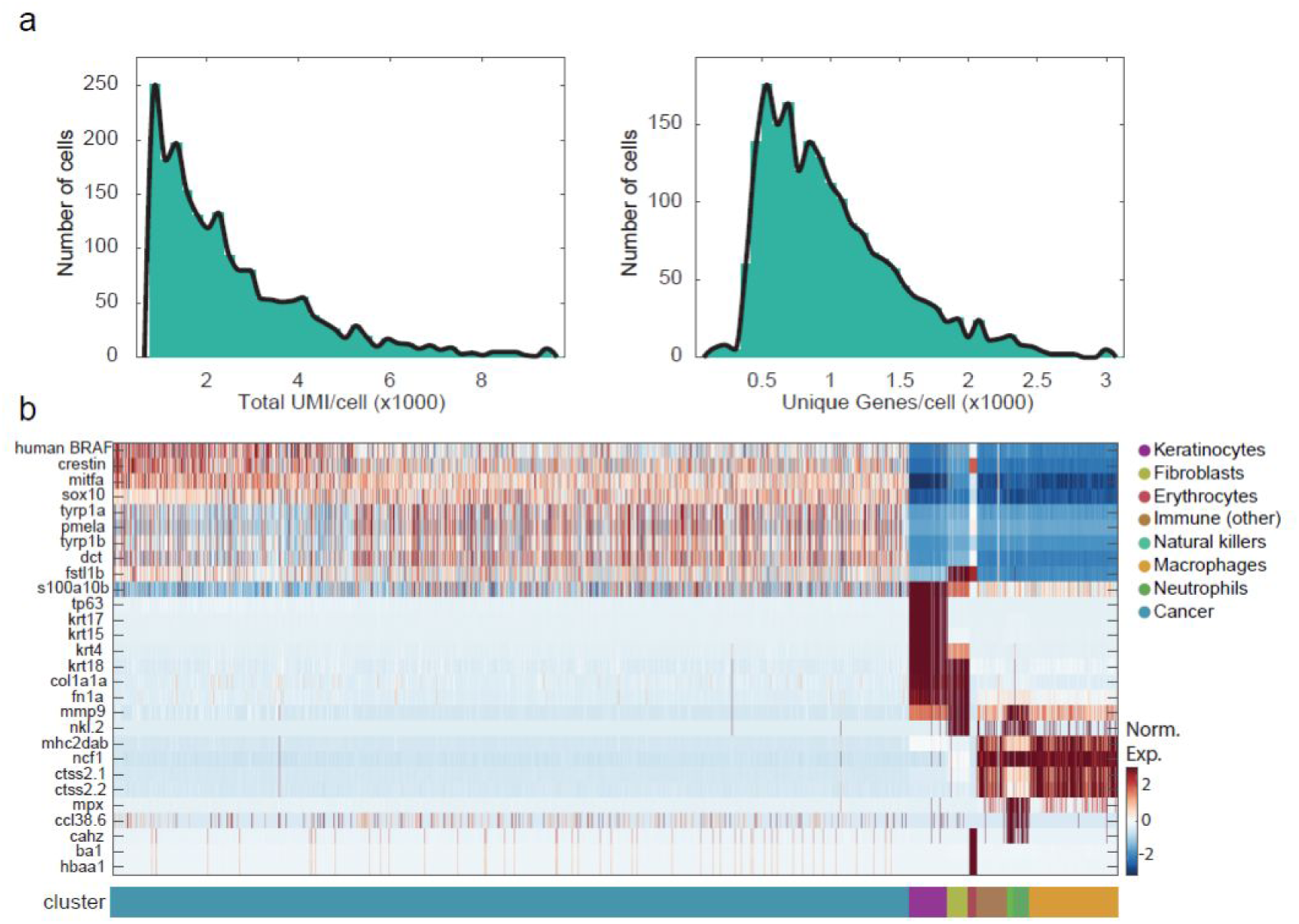
Single-cell RNA-Seq on zebrafish melanoma statistics and clustering. (a) Histogram of the number of transcripts (left panel) and genes (right panel) detected per cell. (b) Heatmap of all cells clustered by hierarchical clustering (see Methods), showing selected marker genes for every population (Table S1). The bottom bar indicates the assigned cell type identity.

**Supplementary Figure 2.**
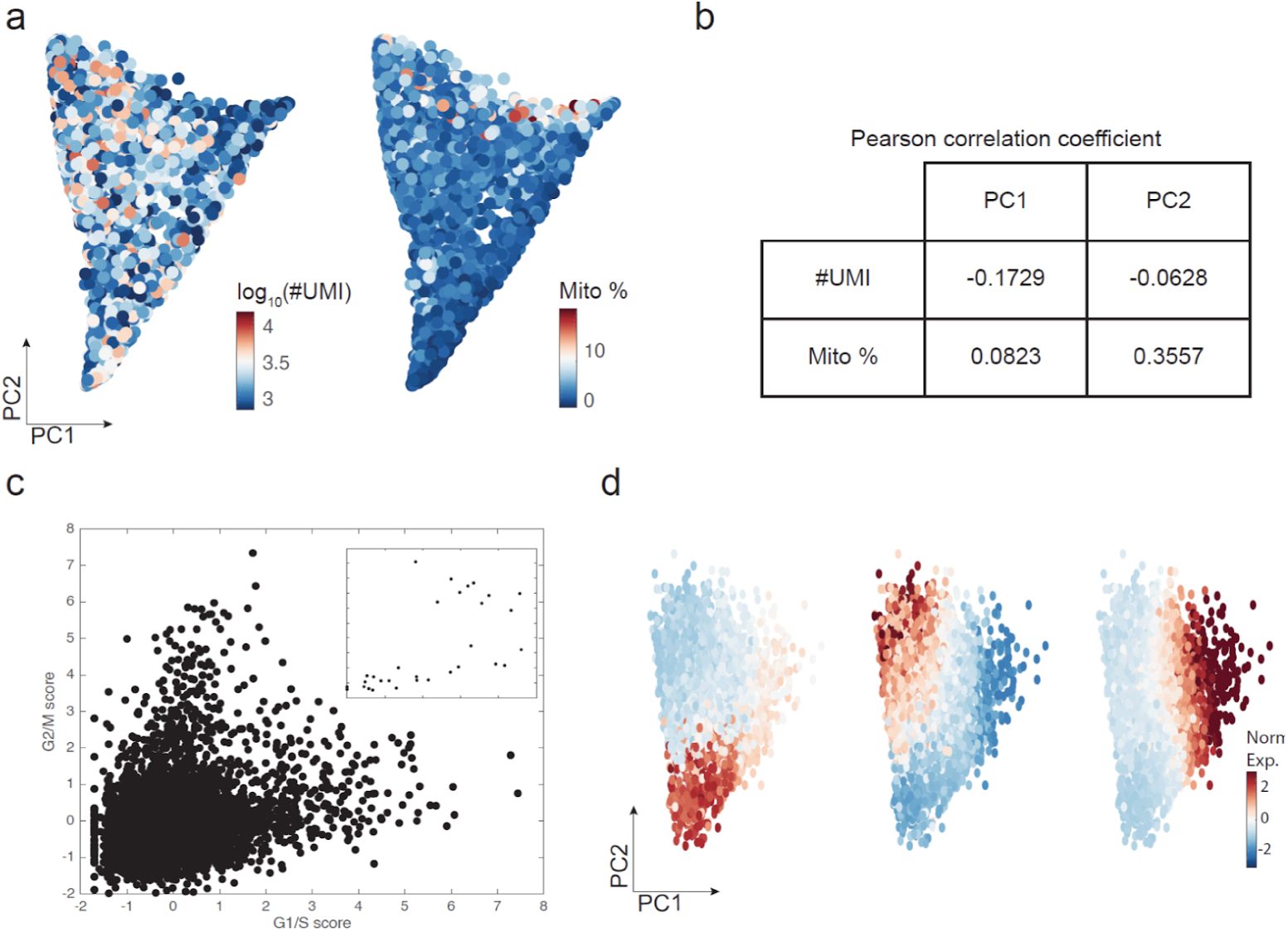
The PCs are not driven by technical aspects or smoothing. (a) PCA of the cancer cells colored by total number of UMI detected (left) and mitochondrial genes percentage (right). (b) Table describing Pearson correlation coefficients with PC1 and PC2 scores. (c) Cell cycle inferred from single-cell RNA-seq. x-axis show the average expression of G1/S genes and y-axis show G2/M genes. There are no clear cycling cells. (d) PCA on non-smoothed cancer cells. Cells are colored by the smoothed expression of genes marking each of the three transcriptional programs: *sox2* for neural crest (left), *dct* for mature melanocyte (middle) and *jun* for stress-like (right).

**Supplementary Figure 3.**
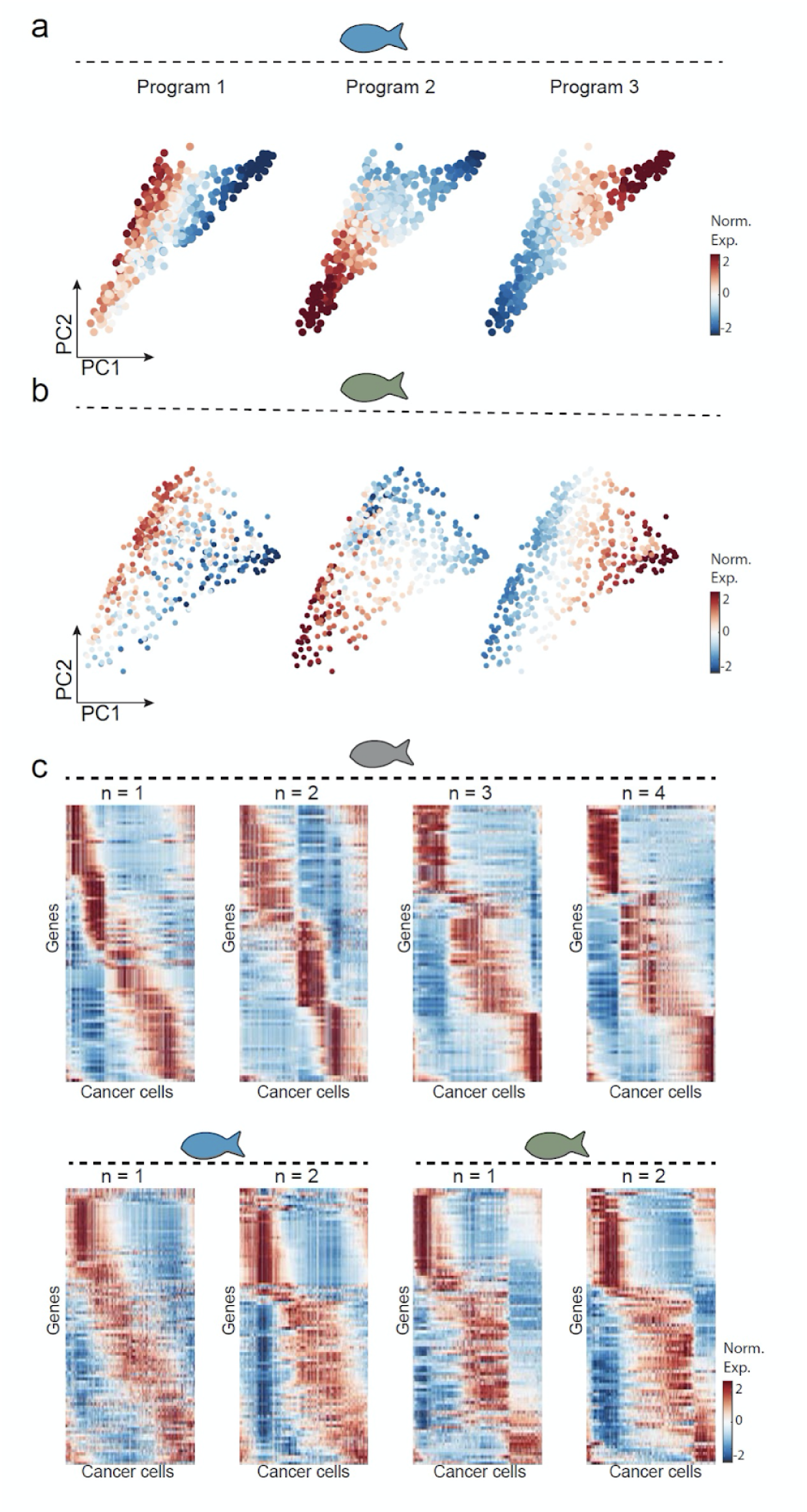
Single-cell RNA-Seq on three zebrafish melanoma tumors. (a,b) Gene expression levels for the neural crest (left), mature melanocyte (middle) and stress-like (right) programs, mapped onto PCA on the tumor cancer cells for each biological replicate. (c) Heatmaps showing the expression of all genes defining the neural crest, mature melanocytes and stress-like transcriptional program, for each biological and technical replicate.

**Supplementary Figure 4.**
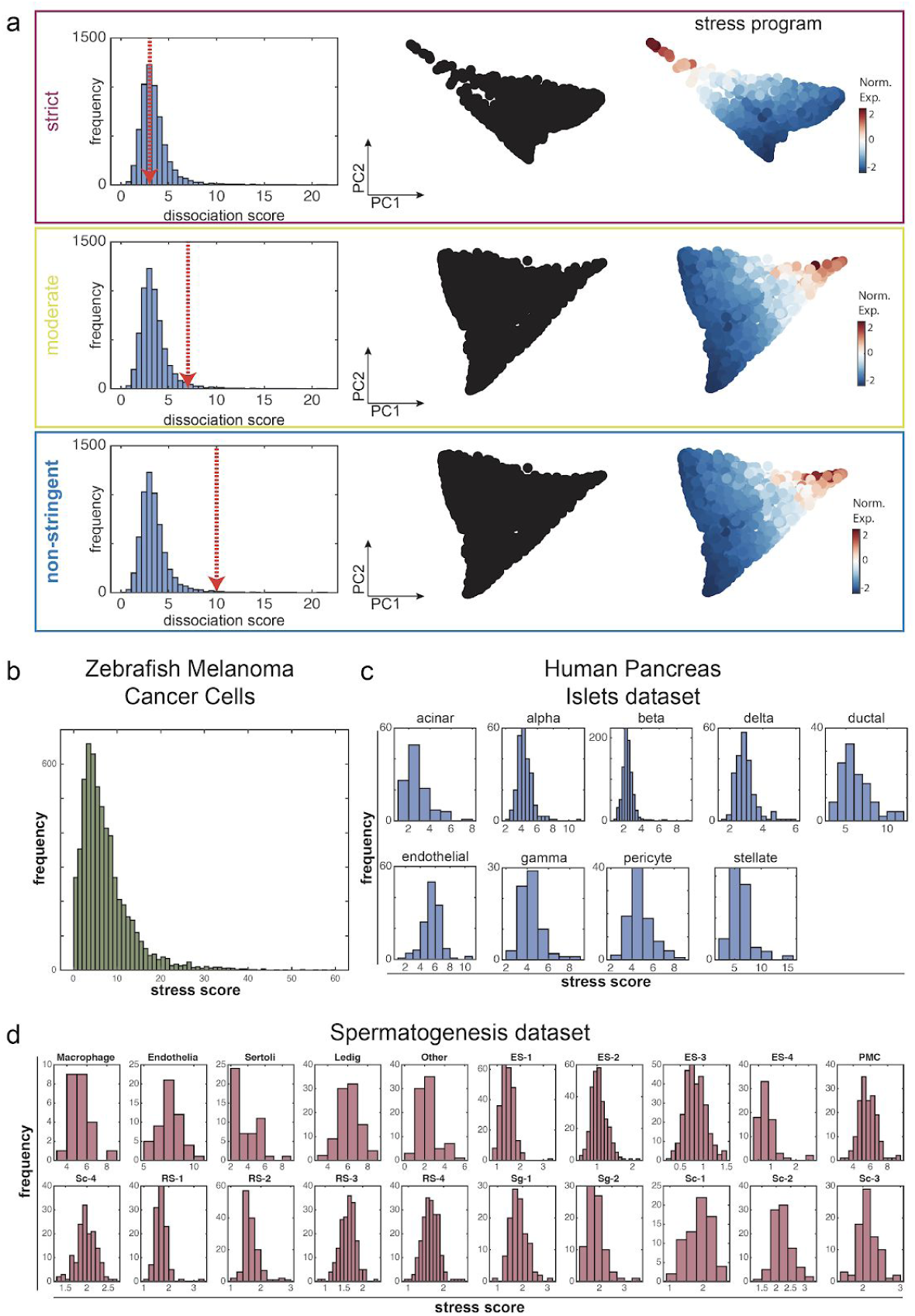
Stress-like gene program is not an artifact of dissociation. (a) *In silico* purification based on Van Den Brink et al. Left - dissociation score distribution across all cancer cells with strict (top), moderate (middle) and non-stringent (bottom) thresholds. Middle – PCA on the cancer cells left after removing all cells that do not meet the dissociation score threshold. Right – stress-like program remains, using all three thresholds. (b) Distribution of stress-like program score, defined as the sum of expression of all genes of this program divided by the expression of all genes in a given cell, across all zebrafish cancer cells. (c,d) Same as (b) for human islets (Baron et al., 2016) and spermatogenesis (Xia et al., 2018) datasets, separated according to cell types.

**Supplementary Figure 5.**
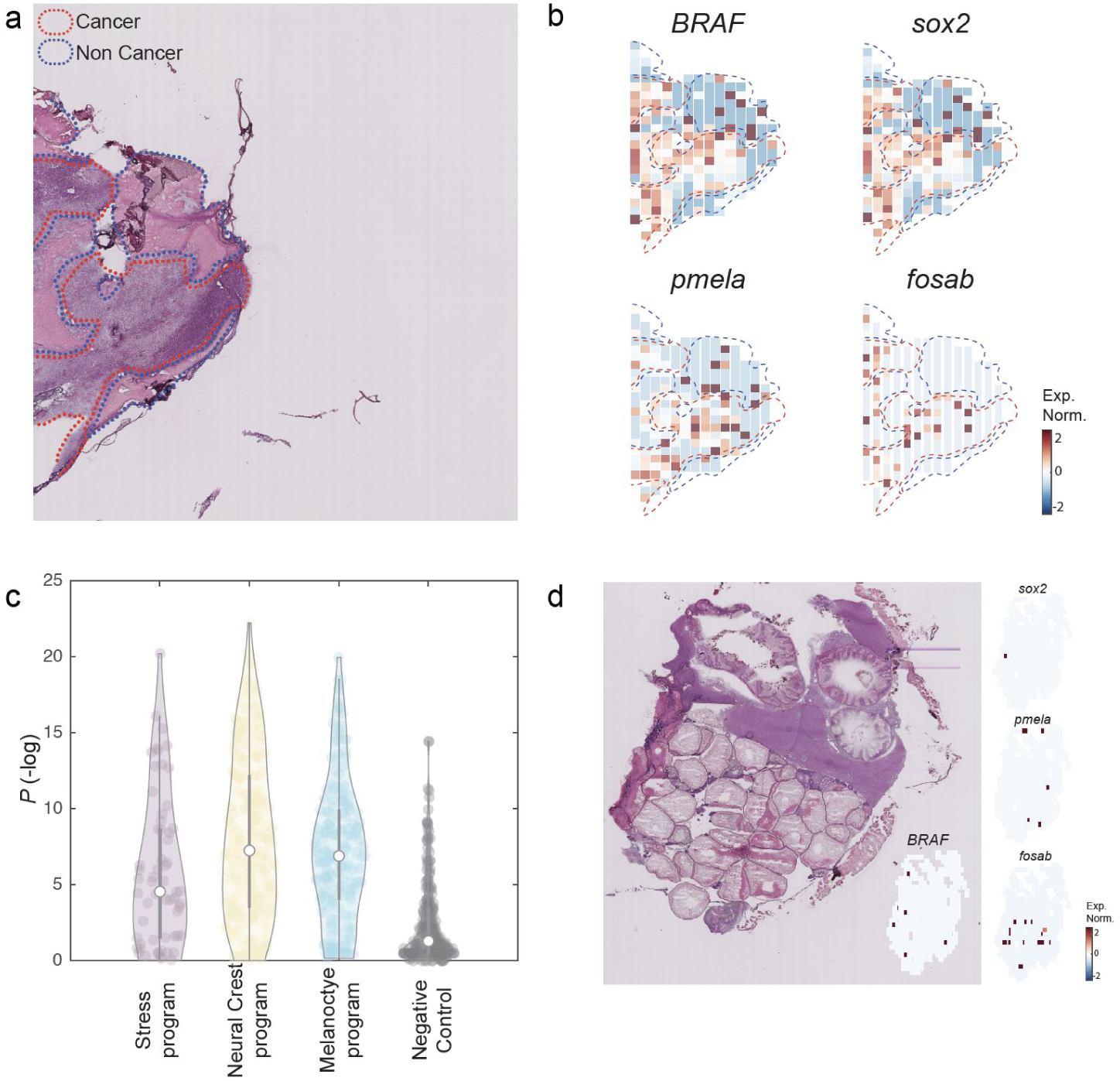
The transcriptional programs involved in the cancer cell states are enriched in cancer areas compared to non-cancer areas in additional section. (a) Hematoxylin and eosin stain of a zebrafish transplanted tumor section. Red and blue dotted lines mark cancer and non cancer areas, respectively. (b) Gene expression profiles of the indicated genes obtained by spatial transcriptomics performed on a section adjacent to the one shown in panel a. (c) Violin plots indicating the enrichment of each gene (Man-Whitney test, −log_10_ of the P-value) in each of the indicated programs. The negative control represents a randomly selected set of 200 genes. (d) Hematoxylin and eosin stain of a non-tumor zebrafish section and gene expression profiles of the indicated genes obtained by spatial transcriptomics performed on an adjacent section.

**Supplementary Figure 6.**
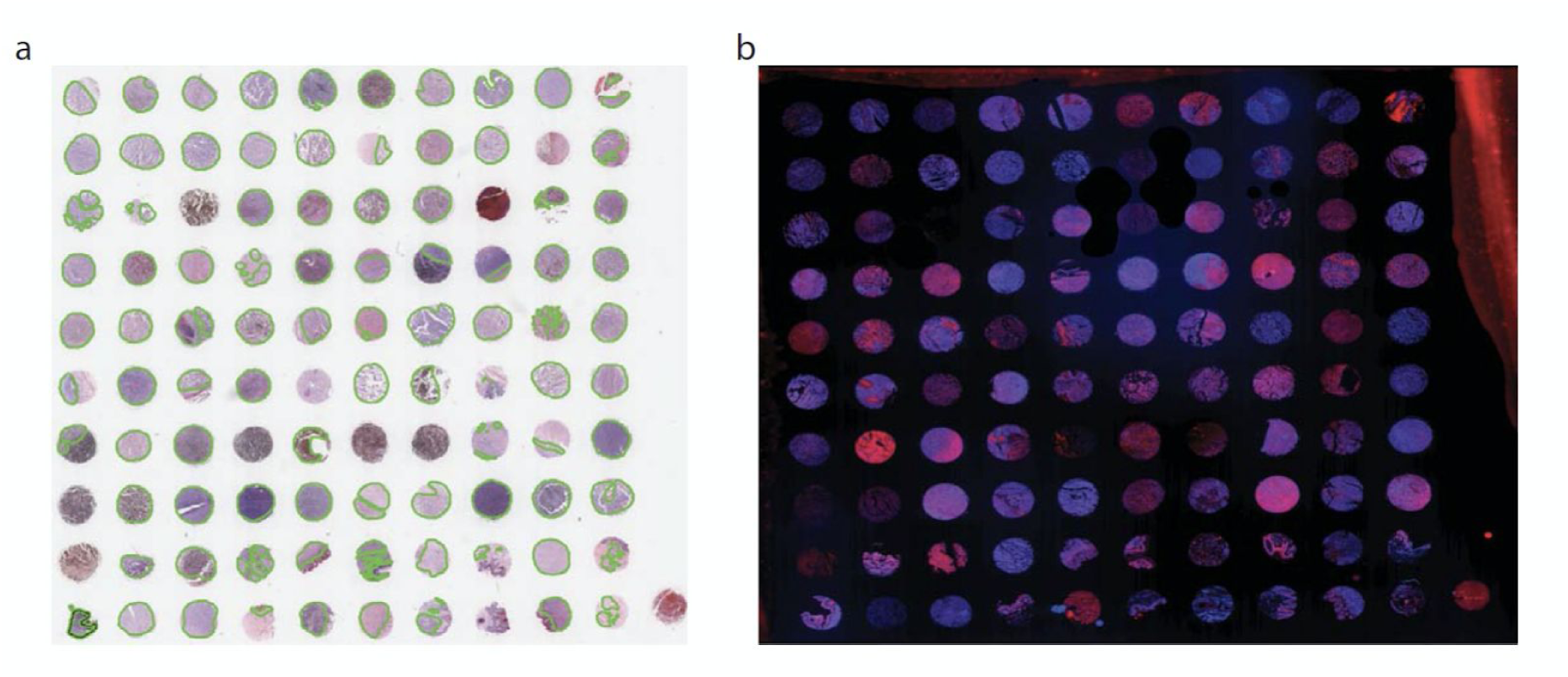
Immunofluorescence of FOS protein, a stress-like program marker, in human melanoma sections. **(a)** H&E staining of a tumor microarray containing 100 human melanoma cores (Biomax, ME1004g). Green lines indicate cancer areas. **(b)** FOS immunofluorescence (red) and DAPI (blue) staining performed on the tumor microarray shown in (a).

**Supplementary Figure 7.**
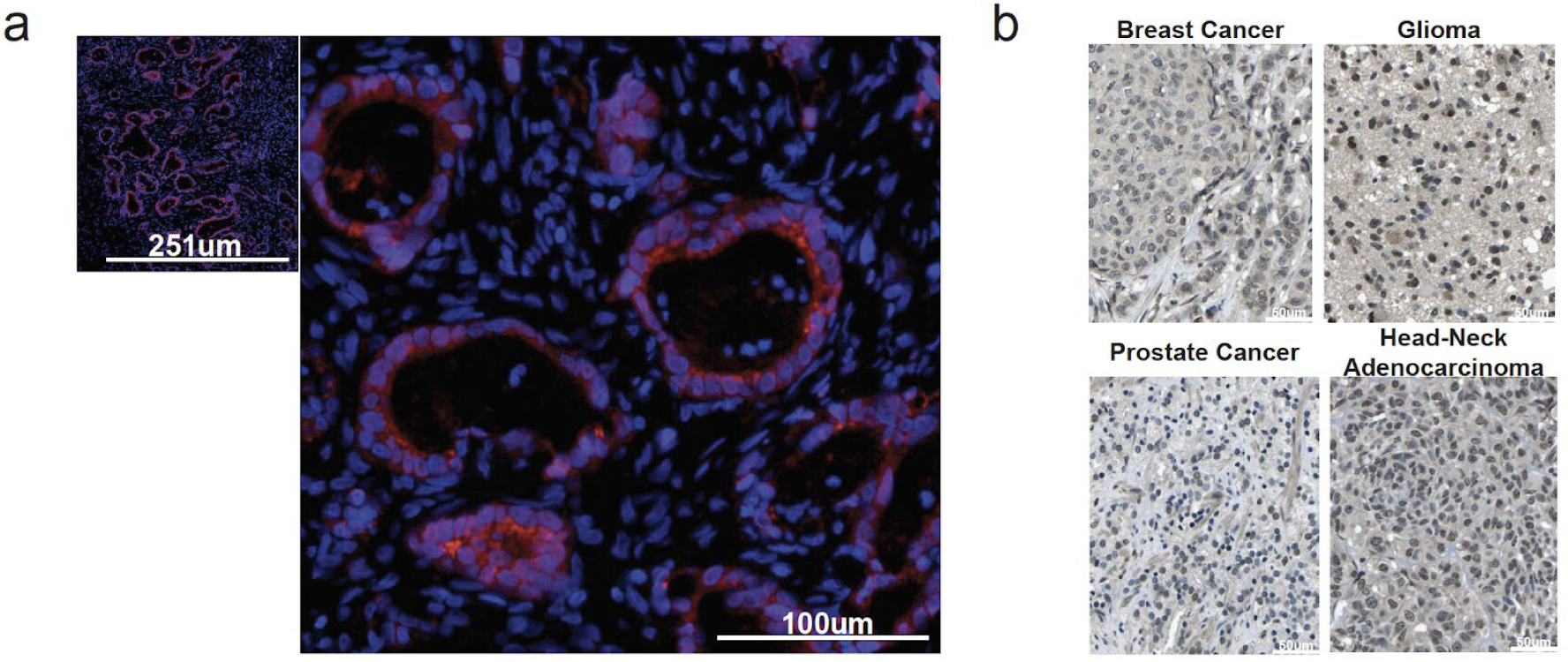
Stress-like cells are conserved at the protein level across other cancer types. (a) FOS immunofluorescence images of pancreatic ductal adenocarcinoma (PDAC). Red indicates FOS expression and blue indicates nuclear stain (DAPI). (b) FOS immunohistochemistry images for the human protein atlas (Thul et al., 2017; Uhlén et al., 2015) for breast cancer, glioma, prostate and head-neck adenocarcinoma cancer.

**Supplementary Figure 8.**
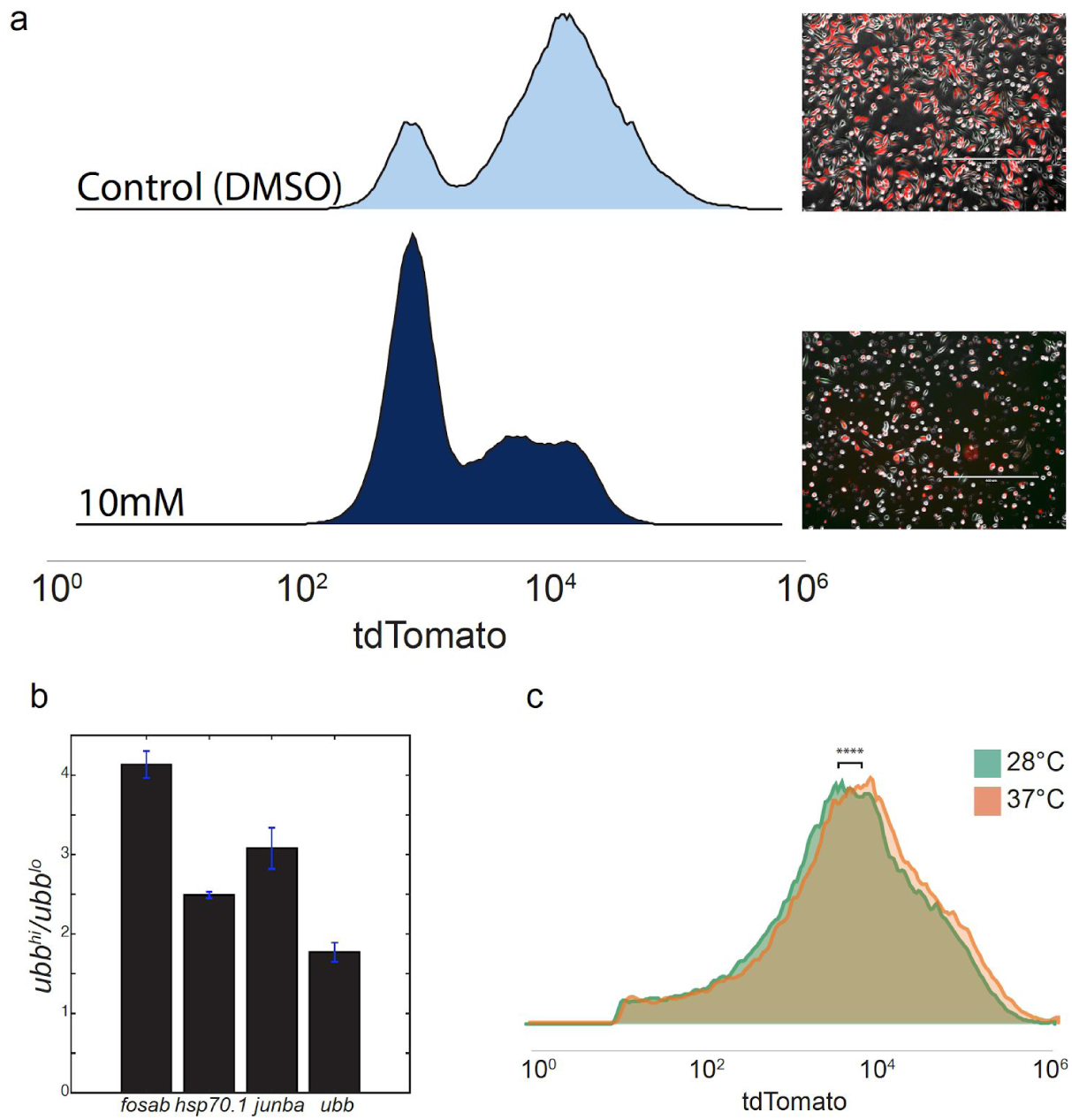
Validation of the *ubb*-tdTomato-NTR-ZMEL1 cell line as the stress-like program. (a) Cells treated with 10mM (lower panel) show change in tdTomato intensity measured by FACS compared to untreated (top panel). Histograms represent distributions of tdTomato intensity of 100,000 events recorded. Images of the cells are attached to each treatment. (b) Bar plot describing the fold change in expression of the indicated genes related to the stress-like program, when comparing high *ubb* cells to low *ubb* cells measured by qPCR. (c) Histograms represent distributions of tdTomato intensity of 100,000 events recorded under optimal growth (green) and heat-shock (orange) conditions. Under heat-shock conditions, cells have higher intensities compared to the optimal conditions.

**Supplementary Figure 9.**
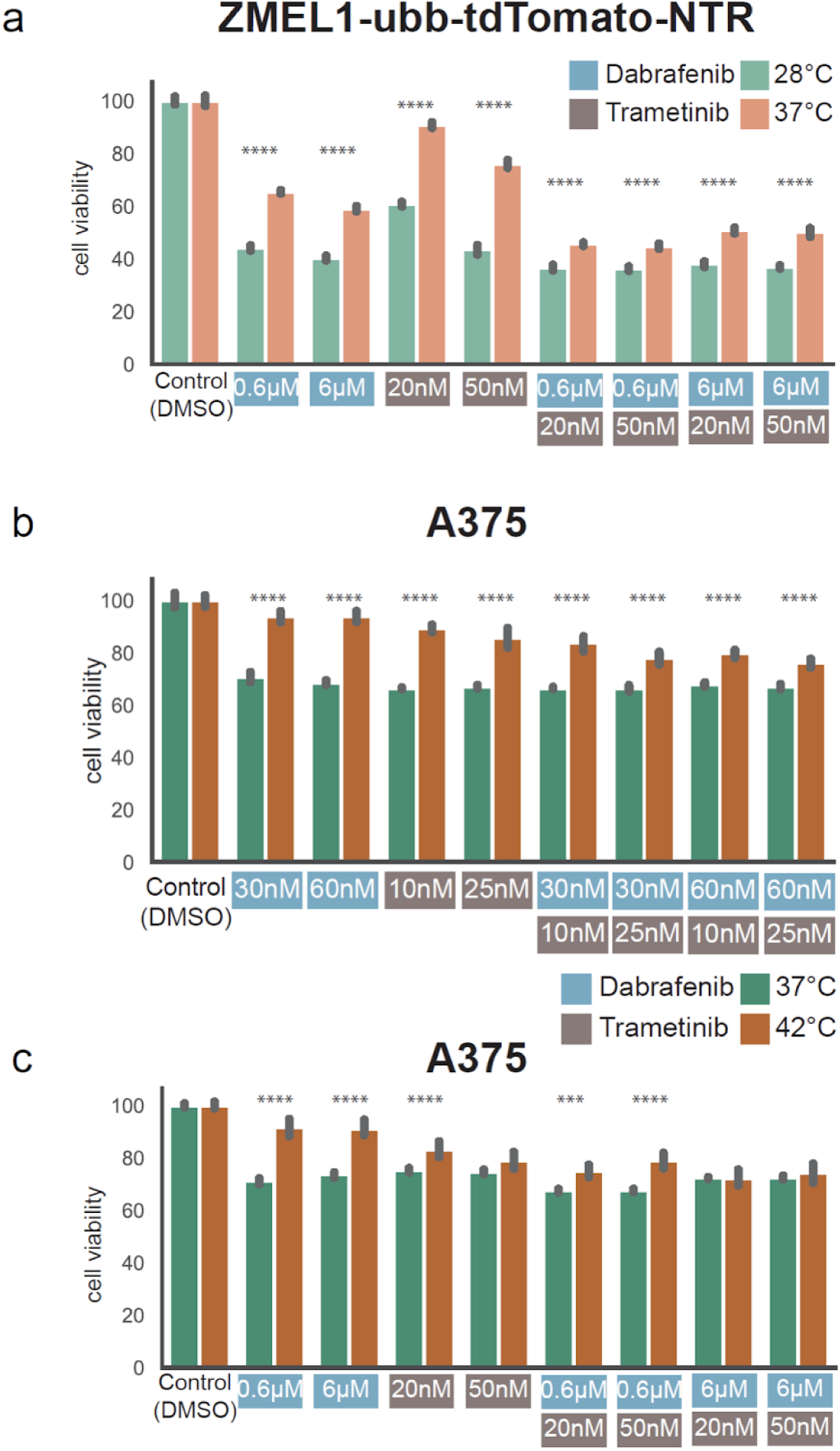
ZMEL1-*ubb*-tdTomato-NTR and A375 cell viability assays under heat shock and drug treatment. (a) Bar plots representing cell viability across culturing conditions (optimal/heat shock) for *ubb*-tdTomato-NTR ZMEL1 in high drug treatments. (b,c) Same as (a) for human melanoma cell line A375 in low and high drug treatments, respectively.

